# Mapping *Pc94*-Mediated Crown Rust Resistance in Oat Reveals a zfBED NLR and Introgression Variation

**DOI:** 10.64898/2026.01.23.701375

**Authors:** Erin L. P. Moreau, Peter J. Maughan, Rebecca Spanner, Eric N. Jellen, Matthew J. Moscou, Shahryar Kianian

## Abstract

Host resistance is a critical component of oat crown rust disease management. *Pc94* is a qualitative resistance locus derived from diploid *Avena strigosa* with several independent introgressions into *A. sativa* that have been used in cultivar deployment. Quantitative trait locus (QTL) analysis combining previously published data for a historic *A. strigosa* population segregating for *Pc94* revealed a large effect QTL on the distal end of *A. strigosa* chromosome 7A. Genome assembly of the parents identified a cluster of five nucleotide binding site leucine-rich repeat receptor (NLR) candidate genes within the QTL region. A single candidate NLR with an integrated zinc finger BED domain, *AstNLR94*, was determined as necessary for *Pc94* resistance based on map-based cloning and forward mutagenesis. A presence/absence allele specific PCR marker was designed in *AstNLR94* and verified for accuracy and specificity in a diverse panel of *A. strigosa* and *A. sativa*. *Pc94* introgressions in *A. sativa* ranged in size from 1.7-71 Mbp and two different introgression locations appear to have occurred. In *A. sativa* ‘Leggett’, a 6.3 Mbp *Pc94* introgression is located at the end of chromosome 2A, and the same sized introgression was discovered in the OT3098 v2 genome. Finally, a QTL analysis identified an additional minor resistance locus on *A*. *strigosa* chromosome 4A, which has complicated previous efforts to characterize the *Pc94* locus. This is the first report of an NLR gene underlying disease resistance in *Avena* spp. and delivers a *Pc94* marker for marker assisted selection to produce disease resistant cultivars.

**Key message:** We mapped a zfBED-NLR encoding gene necessary for *Pc94* resistance, developed a diagnostic marker, and revealed diverse introgression sizes, clarifying *Pc94*’s history and utility for durable oat crown rust resistance.

## Introduction

Common oat (*Avena sativa* L., genome *A_s_A_s_C_s_C_s_D_s_D_s_*) is widely grown in temperate regions for feed, food, forage, cosmetics, and as a cover crop. Annual grain production for 2023 was estimated to be 18 million metric tons, which ranks as 7^th^ highest among the cereals (FAO 2024). Oat is considered a uniquely healthy grain as it contains heart-healthy dietary fiber, antioxidants, and high-quality oil and protein (Butt et al. 2008; Mathews et al. 2020; Peterson 2001). Regular oat consumption is well documented to reduce the risk of diabetes and cardiovascular disease (Daou and Zhang 2012; Mathews et al. 2020).

Oat crown rust (OCR) caused by *Puccinia coronata* f. sp. *avenae* P. Syd. & Syd (*Pca*) is the most economically important oat disease worldwide (Nazareno et al. 2018). The biotrophic disease reduces yield and quality of grain and forage and impacts winter hardiness of winter oat (Doehlert et al. 2001; Fulcher et al. 2020; Lovatto et al. 2021; Simons 1985). *Pca* reproduces asexually on oat, with repeated two-week cycles of urediniospore infection and sporulation leading to OCR epidemics under favorable environmental conditions (Simons 1970). In severe cases, grain yield losses of 35-50% have been documented in major oat producing regions (USDA-ARS CDL 2014). Alternate hosts in the genus *Rhamnus*, including the widespread common buckthorn (*R. carthartica),* facilitate sexual reproduction of *Pca* in Europe, the Middle East, North America, and to a limited extent in Australia (Dietz 1926; Henningsen et al. 2024a; Simons 1970), giving rise to highly adaptable, genetically diverse *Pca* populations (Hewitt et al. 2023; Miller et al. 2021). Somatic hybridization also plays a role in generating genetic variability and shifting virulence (Henningsen et al. 2024a).

At the molecular level, the interaction between *Pca* and oat can be understood through the lens of the zigzag model (Jones and Dangl 2006). As an adapted pathogen of oat, *Pca* effectively colonizes oat through suppression of defense mechanisms through the action of secreted effector proteins. Oat has evolved an arsenal of Resistance (R) proteins that recognize effector proteins directly or indirectly and activate a resistance response. Resistance is often mediated through a hypersensitive response that is generally associated with cell death, which in turn restricts pathogen development and spread. However, *Pca* strains or races with mutations rendering the compromised effector non-functional or unrecognizable by oat can evade *R* gene detection and cause disease. An evolutionary arms race follows, driving strong diversifying selection on both pathogen effectors and plant *R* genes (Ma and Guttman 2008; Plissonneau et al. 2017; Tamborski and Krasileva 2020).

Qualitative (race specific) *R* genes are routinely used in breeding and have been extensively studied, with most cloned *R* genes encoding intracellular immune receptors belonging to the nucleotide binding, leucine rich repeat (NLR) family (Jones et al. 2024). These proteins have highly variable, modular structures consisting of the hallmark nucleotide binding (NB) domain and leucine-rich repeats (LRRs) domains. Many NLRs also contain an N-terminal coiled coil (CC) domain or a Toll/interleukin-1 receptor (TIR) domain. About 10% of NLRs have additional non-canonical Integrated Domains (NLR-IDs), such as WRKY domains or zinc finger BED (zfBED) domains, which may serve as effector decoys (Kroj et al. 2016; Marchal et al. 2022; Sarris et al. 2016).

Host resistance is a cornerstone of OCR disease management. Over 90 *Pc* loci conferring qualitative resistance have been described, although some of these gene designations are likely redundant or describe allelic variants (Hewitt et al. 2023; Miller et al. 2021; Park et al. 2022). Effective qualitative resistance from *A. sativa* sources were largely exhausted by the mid 1900’s, leading to searches for additional genes in closely related oat species (Fleischmann and Baker 1971; Simons and Murphy 1955). Most named *Pc* loci used in modern oat breeding originated from wild hexaploid *Avena sterilis* (genome *A_s_A_s_C_s_C_s_D_s_D_s_*), wild tetraploid *Avena magna* (*C_m_C_m_D_m_D_m_*), cultivated tetraploid *Avena abyssinica* (*A_a_A_a_BB*), and cultivated diploid *Avena strigosa* (*A_a_A_a_*) (Park et al. 2022). Due to *Pca*’s ability to rapidly adapt and acquire virulence to deployed qualitative *R* genes (Carson 2009; Chong and Kolmer 1993; Moreau et al. 2024), intentional stacking of multiple qualitative *R* genes with more durable quantitative resistance holds the greatest promise for enduring disease resistance. Efficient marker assisted selection requires high quality and high throughput markers, but their development in oat has lagged behind that of other cereal crops. Recent advances in long read sequencing have enabled chromosome-scale genome assemblies for hexaploid oat (Avni et al. 2025; Kamal et al. 2022; Peng et al. 2022; PepsiCo 2021; Zhang et al. 2025), accelerating *R* gene mapping and marker development (Abdullah et al. 2023; Admassu-Yimer et al. 2022; Argenta et al. 2025; Nguyen et al. 2025; Toporowska et al. 2021; Wight et al. 2024). However, no *Pc* locus has been resolved to a candidate gene or cloned to date.

*Pc94* is a qualitative *Pc* locus derived from cultivated diploid *A. strigosa* and deployed in spring oat cultivars for the Northern United States and Canada. It was introgressed into *A. sativa* in two separate instances (Aung et al. 1996; Rines et al. 2007) before the two donor lines’ resistances were determined to be controlled by the same major locus (Park et al. 2022; Rines et al. 2007). There have been conflicting reports as to whether *Pc94* resistance is mediated by a single gene (Rines et al. 2007; Simons et al. 1959) or five tightly linked ones (Wise et al. 1996; Yu and Wise 2000). The first introgression is the source of *Pc94* in most modern cultivars (Caffe-Treml 2021a, b, c, d, 2023; Mitchell Fetch et al. 2007; Smith 2020). The second introgression is the source of *Pc94* in ‘ND Carson’ (McMullen 2025).

*Pc94* has shown relatively long durability in the field, although virulence to the locus has recently increased. In Canada, where *Pc94* was first released in ‘Leggett’ in 2007, virulence was almost undetected in *Pca* surveys from 1993-2015 (Chong 2000; Chong et al. 2008; Menzies et al. 2019). Since 2020, virulence to *Pc94* increased to 20-28% of isolates collected from commercial fields in the Eastern Prairie Region (Menzies et al. 2024a; Menzies et al. 2023; Menzies et al. 2024b; Menzies et al. 2021; Menzies et al. 2022). In the USDA virulence survey, only 0-11% of isolates were virulent to *Pc94* from 2005-2021 (Moreau et al. 2024). *Pc94* began to break down in the Northern United States starting in 2022, with 52-83% of isolates virulent in recent surveys (Moreau et al. 2025). In contrast, virulence to *Pc94* has remained low in Southern United States isolates at 5-10%, Poland (∼8-15%) (Paczos-Grzęda and Sowa 2019; Sowa and Paczos-Grzęda 2020), and in Australia (4%), where it was recently added to virulence monitoring (Henningsen et al. 2024b). To our knowledge, *Pc94* is not currently being used in breeding or virulence monitoring in other regions such as South America or Western and Northern Europe (Dietz et al. 2024).

Despite rising virulence to *Pc94* in the Northern United States and Canada, fine-mapping the locus and generating a linked marker are important for deploying *Pc94* in combination with other resistance genes into new regions and for reliable genotyping when its phenotype is suppressed by *Pc38*. A marker for *Pc94* was previously developed (Chen et al. 2007; Chong et al. 2004) but is not being used because it gives false negative results in known *Pc94* carriers such as Leggett.

Here we report *AstNLR94*, an NLR with an integrated zfBED domain necessary for *Pc94* resistance originally derived from *A. strigosa* chromosome 7A. We developed a high throughput *Pc94* PACE marker which identifies *AstNLR94* in both *A. strigosa* and *A. sativa*. We investigated the size and location of the introgressions containing *Pc94* in *A. sativa* cultivars and trace the introgression through pedigrees. This is the first report of a fine-mapped *R* gene in *Avena*. Additionally, we identified a minor segregating *R* locus in the *Pc94* donor background via QTL analysis which was not transferred into *A. sativa* but has complicated previous *Pc94* mapping efforts in *A. strigosa*.

## Materials and Methods

### Germplasm

Seed for the mapping population parents CIav 3815 (PI 131641, *Pc94* carrier) and CIav 1994 (PI 53626, OCR susceptible) were provided by the late Dr. Dave Hoffman (USDA-ARS) to Dr. Rick Jellen. Accessions CIav 3815 and CIav 1994 are the parents of the historical recombinant inbred line (RIL) population used to map *Pc81-85* (syn. *Pc94*). CIav 3815 originated from Krakow, Poland, and was indexed in the USDA National Plant Germplasm System (NPGS) in 1939. CIav 1994 is recorded originally as a wild *A_a_* diploid *Avena wiestii* collected from Al Jīzah, Egypt, however recent genetic analysis has shown this accession is *A. strigosa* (Maughan et al. 2019). The CIav 1994 used in this study was determined to be the same as CIav 1994 sourced directly from the NPGS in 2019 using 45 polymorphic *A. strigosa* SNPs from the USDASoyWheOatBar-3 K Illumina iSelect array (Jason Fiedler, personal communication, Supplementary Table 1). CIav 3815 and CIav 1994 were crossed to generate 7 F_1_ plants which were selfed for two generations to produce F_2:3_ families.

### Genome assembly and annotation

We generated two high quality, annotated genomes for the two parental accessions using PacBio HiFi long read technology coupled with Hi-C (chromatin contact conformation) data. High molecular weight (HMW) DNA was extracted from fresh leaf tissue for each accession using a CTAB-Genomic-tip protocol (Vaillancourt and Buell 2019). DNA was quantified using a spectrophotometer and assessed for quality, including concentration (>800 ng/mL) and purity (260/280 and 260/230 ratios ≈ 2.0). For PacBio HiFi sequencing, HMW DNA was sheared to 17 kb using a Diagenode Megaruptor (Denville, NJ) and converted into SMRTbell libraries with the SMRTbell Express Template Prep Kit 2.0 (Pacific Biosciences, Menlo Park, CA). Libraries were size-selected (>10 kb) on a Sage Science BluePippin (Beverly, MA). Sequencing was performed at the Brigham Young University DNA Sequencing Center (Provo, UT, USA) on the PacBio Revio platform using the Sequel II Sequencing Kit 2.0 with Sequencing Primer v5 (PN:102-067-400; PacBio, London, UK) and the Sequel Binding Kit 2.2, run for 30 h with adaptive loading following PacBio SMRT Link recommendations.

Primary contig assemblies were generated using hifiasm v0.19.8 (Cheng et al. 2021) with default parameters for an inbred species (−l0) while scaffolding of the primary contig assemblies into pseudo-molecules was accomplished using Hi-C data generated for each accession. Hi-C libraries were generated by Phase Genomics (Seattle, Washington, USA) from fresh leaf tissue at an industry-standard depth of ∼100 million read pairs per Gb of genome size. Hi-C reads were aligned to their respective primary contig assemblies using the Arima-HiC pipeline (A160156_v03, Arima Genomics, Carlsbad, California, USA). Only uniquely mapped read pairs were retained for scaffolding. Contigs were clustered, ordered, and oriented using YaHS (Zhou et al. 2023), a proximity-guided scaffolder, followed by manual inspection and correction using JuiceBox (Durand et al. 2016a). Inter-contig gaps within scaffolds were padded with 100 Ns. Genome quality value (QV) was estimated using Merqury (Rhie et al. 2020) by comparing k-mer spectra derived from the HiFi sequencing reads to the assembled genomes to quantify base-level consensus accuracy.

Prior to de novo gene annotation, repetitive elements were identified and masked using RepeatModeler2 v2.0.1 (Flynn et al. 2020) and RepeatMasker v.4.1.2 (Smit et al. 2013–2015). Each soft-masked genome was annotated with Braker3 v.3.0.7 (Gabriel et al. 2024) using a combined IsoSeq dataset consisting of the raw IsoSeq data from all *A. sativa* accessions reported by the PanOat consortium project (Avni et al. 2025), RNAseq data derived for both accessions (see RNASeq section below), as well as the publicly available *A. strigosa* RNAseq data generated by Li et al. (2021) as primary evidence, with the IsoSeq being considered the reference (i.e., if two gene models overlapped, the Isoseq model was deemed more correct). The Viridiplantae OrthoDB v.11 (Kuznetsov et al. 2023), Poales UniProtKB (www.uniprot.org/uniprotkb), and the NCBI Poales RefSeq (www.ncbi.nlm.nih.gov/protein) databases served as training evidence for AUGUSTUS. Gene models without a canonical start and stop codon were removed, as well as any models where the length of the gene was less than 150bp (protein < 50AA). Furthermore, any models that had significant matches to the transposable element database (TREP; https://trep-db.uzh.ch/) were also removed. Lastly, we used InterProScan (*e*-value < 1.0e-6) (Jones et al. 2014) and the UniProt-Swiss databases (downloaded 7/2023) to add gene ontology (PANTHER-15.0, Pfam-33.1, PIRSF-3.10, PIRSR-2021_02, PRINTS-42.0, and SUPERFAMILY-1.75) and functional description to the remaining gene models. Chromosomes and gene models were named as described by Jellen et al. (2024). The completeness of the gene space in the genomes and annotations for each accession was evaluated with BUSCO v5 (Manni et al. 2021) using the poales_odb10 protein dataset (Supplementary Fig. 1). Genome assembly statistics are available in Supplementary Table 2.

### Preliminary QTL analysis

We compared linkage groups (LGs) of the high density linkage maps constructed for the CIav 3815 x CIav 1994 RIL population (Latta et al. 2019) to the assembled genomes and reorientated LGs to match the International Oat Nomenclature Committee chromosome nomenclature: LG 4 became LG 5 with inverse orientation, LG 5 became LG 4, and LG 6 was inversed. Supplementary linkage map data were integrated including roughly binned RFLP markers and the *Pc94* locus from the previous map (Kremer et al. 2001). In the new map, the *Pc94* locus (labeled after *Pca* isolates I202, I263, I290, PC54, PC62) was binned with distal markers Oat18480.1, Oat38313.1, Oat408.1, Oat5006.1, and Oat18466.1 on LG7 at 136.1 cM. To verify the position of *Pc94*, a preliminary QTL analysis for the RIL population was performed using the 2019 linkage map with LG updated order (Supplementary Table 3) and the original phenotypic data from *Pca* isolates I202, I263, I290, PC54, PC62 (Wise et al. 1996) in R Statistical Software v4.4.1 with r/qtl2 v0.36 (Broman et al. 2019; R Core Team 2022). Genotype probabilities were calculated with the function calc_genoprob and a genome scan for a single-QTL binary model was performed with scan1 with default parameters. For each isolate, 1,000 permutations were performed to determine significance thresholds at α=0.05. Sequence tags for Oat18480.1, Oat38313.1, Oat408.1, Oat5006.1, and Oat18466.1 were blasted against the CIav 3815 genome assembly to determine the approximate area to screen for F_2_ recombinants (Camacho et al. 2009). PACE markers were designed based on single nucleotide polymorphic sites to saturate the region.

### Recombinant screening

We screened F_2_ plants (n=851) for recombinants encompassing the *Pc94* locus. DNA was extracted using an SDS method modified from Edae et al. (2024). Briefly, young leaf tissue was harvested into 96-well plates and freeze dried overnight in a lyophilizer. Samples were ground for 2 min at 20 rpm using a Genogrinder (SPEX Sample Prep) and 4 mm stainless steel grinding balls (Cole-Parmer). 300 µl extraction buffer (200 mM TrisHCL pH 8.0, 250 mM NaCl, 25 mM EDTA, 0.5% SDS, ddH20) was added to each well, and the plates were gently shaken before addition of 300 µl of chloroform:isoamyl alcohol (24:1). The suspension was mixed for 3 min on a plate shaker followed by centrifuging for 20 min at 4000 rpm, and the supernatant (150-200 µl) was transferred to a new plate. 300 µl cold isopropanol was added to each well of supernatant, the plates were mixed through gentle shaking, then incubated on ice for 15 min. Plates were centrifuged for 20 min at 4000 rpm. The isopropanol was decanted and 300 µl 70% ethanol was added to each well then plates inverted to thoroughly wash the pellet. The plates were centrifuged for 20 min at 4000 rpm, the ethanol was decanted, and the DNA pellets were air-dried in a dryer overnight. Then 300 µl of 1xTE low EDTA buffer (10 mM TrisHCL pH 8.0, 0.1 mM EDTA pH 8.0, ddH20) was added to each tube and pellets resuspended. DNA was quantified using a NanoDrop 1000 spectrophotometer (Thermo Fisher, Waltham, MA, United States) and diluted to 15 ng/µl for PACE genotyping.

PACE reactions consisted of 8 µl total reaction volume per sample in H_2_O with 15 ng sample DNA, 4 µl 2x PACE buffer, and 0.1 µl assay mix (12 µM forward primer 1, 12 µM forward primer 2, and 30 µM reverse primer in H_2_O). Each 96 well plate run contained three non-template controls. The cycling conditions on the Lightcycler 480 II (Roche) were preincubation at 94°C for 15 min, 10 touchdown cycles (94°C for 20 sec, 61°C-0.6°C/cycle for 1 min), 30 amplification cycles (94°C for 20 sec, 55°C for 1 min), and endpoint data collection (37°C for 1 min, data acquisition).

If markers needed additional cycles to tighten clustering, an additional three-step recycle was performed with the following parameters: 3 amplification cycles (94°C for 20 sec, 57°C for 1 min) and endpoint data collection (37°C for 1 min, data acquisition).

### Phenotypic screening

We screened recombinant F_2:3_ families using seedling inoculations and three *Pca* isolates, 12SD80 (two replications), 15FL1-2, 15OH12-3. These isolates were chosen because they displayed slightly different infection types from each other in preliminary tests on the original recombinant RILs from the study proposing five tightly linked loci in the *Pc94* region (*Pc81-85*) (Wise et al. 1996).

For inoculation, *Pca* urediniospores were heat shocked at 40°C for 10 min and suspended in mineral oil (Soltrol 170). Fifteen to twenty nine-day-old seedlings per F_2:3_ family were inoculated with spore suspensions at a rate of 4 mg spores/750 ml of mineral oil per inoculation batch (18 F_2:3_ families) using a custom fabricated inoculator and 5 psi air pressure. After the mineral oil dried, plants were placed in the mist chamber overnight with intermittent misting for 40 sec on and 4 min off at 20 to 22°C. Plants were maintained in a greenhouse (21 to 24°C) and fertilized with 5 g of 20-20-20 NPK water soluble fertilizer per liter of water at seedling emergence, 1 day post inoculation (dpi), and 8 dpi. Plants were rated for crown rust reaction at 11-14 dpi.

Seedlings were rated using a modified 0-4 infection type (IT) scale (Murphy 1935; Nazareno et al. 2018), and F_2:3_ families were classified as resistant (all 0-2n), susceptible (all 3-4), or segregating (mixtures of R and S). These values were compared to the PACE results to narrow down the *Pc94* locus to PACE markers P_As_26 and P_As_07. To further narrow down the region, the population was expanded to an additional 860 F_2_ plants and 340 F_4_ plants for recombination screening in the refined region. F_2_ lines with recombination in critical intervals were advanced to the F_3_ generation and screened via PACE markers for homozygosity to fix the recombination events.

### RNAseq

RNA was extracted from the second emerged leaves of CIav 3815, CIav 1994 and homozygous critical recombinants F_3_-233 and F_3_-474 using RNeasy Plant Mini kit (Qiagen). The standard protocol was followed with the additional on-column DNase Digestion. The University of Minnesota Genomics Center performed RNA sequencing preparation, multiplexing, and sequencing on the NovaSeq 6000 or NovaSeq X-Plus (Illumina) targeting 40 million read sequencing depth per sample with 150 bp paired end reads.

Reads were preprocessed using trimmomatic (v0.38) (Bolger et al. 2014) with the following flags: TruSeq3-PE.fa:2:30:10 LEADING:5 TRAILING:5 SLIDINGWINDOW:4:15 MINLEN:36. For the parents and homozygous recombinants, paired reads were aligned with HISAT2 (Kim et al. 2019) to a 300 kb region encompassing the refined QTL interval in CIav 3815. SAMtools (v1.17) (Danecek et al. 2021) was used for conversion to sorted bam files for viewing.

### PCR and sequencing of candidate genes

To test for the presence of candidate gene *G37621* (AVATL.20003a.r1.7AG00000037621) in homozygous critical recombinant F_3_-1239, we PCR amplified *G37621* (primers in Supplementary Table 4) with PrimeStar GXL DNA polymerase (Takara Bio USA) and the following cycling conditions: 98°C for 2 min, 10 touchdown cycles (98°C for 10 sec, 72°C-1°C/cycle for 15 sec, 68°C for 5 min) and 25 amplification cycles (98°C for 10 sec, 62°C for 15 sec, 68°C for 5 min). PCR products were run and imaged on a 1.5% agarose gel with GelRed Nucleic Acid Stain, purified with Monarch PCR and DNA cleanup kit (New England Biolabs), and verified via Oxford Nanopore sequencing (Eurofins Genomics).

### EMS mutagenesis screen

We treated the *A. sativa* ‘*Pc94*’ differential line with ethyl methanesulfonate (EMS) using a protocol adapted from barley (Caldwell et al. 2004; Dinh et al. 2022). Briefly, 175 g of seed (∼5,000 seed) were imbibed with water for 4 hours. The water was replaced with 350 ml 60 mM EMS solution. The EMS rate was determined by preliminary experiments to target 50% *M*_1_ lethality. The solution was shaken at ambient temperature and 100 rpm for 16 hours. The EMS solution was removed, and the seeds were treated with two changes of 350 ml 100mM sodium thiosulfate solution to deactivate the EMS. Seeds were treated to five water washes with shaking for five min each wash. Seeds were transferred to trays covered with filter paper to dry overnight before sowing directly in the field. A total of 739 individual *M*_1_ panicles were collected from the field grown plants. Seeds from each panicle were sown into individual 4-inch pots and screened for loss of resistance to *Pca* isolate 12SD80 with the inoculation procedure described above for a total of 7,507 *M*_2_ seedlings screened. In every inoculation batch, “Marvellous” served as a positive control while the non-mutagenized *A. sativa Pc94* differential served as a negative control. All susceptible *M*_2_ plants were transplanted for DNA extraction and seed increase via selfing. For each susceptible plant, *G37621*, *G37622* (AVATL.20003a.r1.7AG00000037622), and *G37624* (AVATL.20003a.r1.7AG00000037624) were PCR amplified, purified, and sequenced as described above. Six seedlings each of the *G37624 M*_3_ mutants and controls (Marvellous and *Pc94*-WT) were inoculated with genetically and pathologically diverse *Pc94* avirulent isolates 90MN5B-1, 90TX52-1, 90AR100-1, 12NC29, 12SD18, 15FL1-2, 15MS7-1, 15MS16-3, and 15MN23-1 (Miller et al. 2021) to test for isolate specificity in mutant loss of resistance. Original inoculation photos featured in Figure 1C and Supplementary Figure 4 are available on figshare (10.6084/m9.figshare.31038247).

**Fig. 1.**
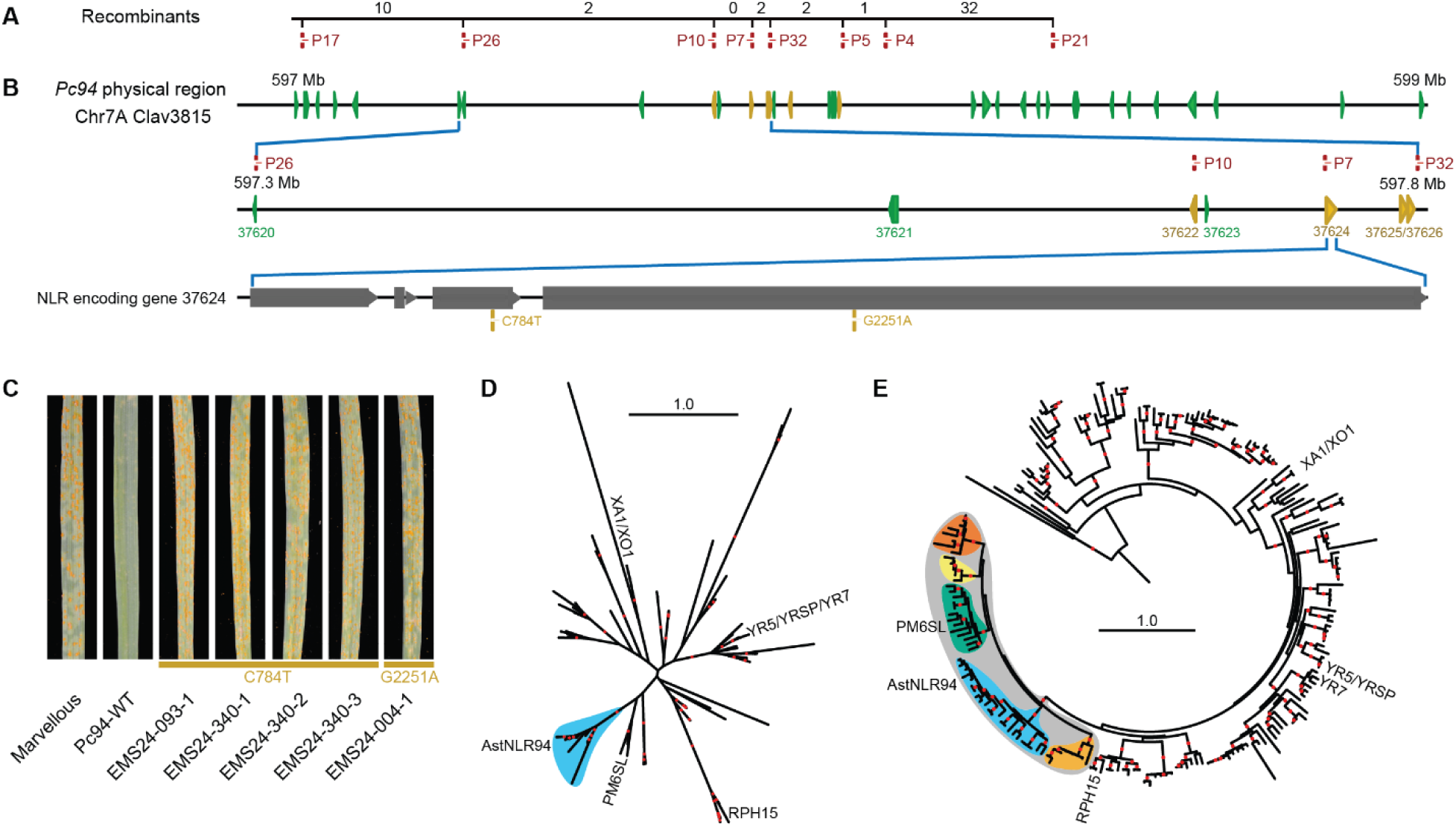
Fine-mapping and mutagenesis of *Pc94*. A. Genetic and physical regions encompassing the *Pc94* region on CIav 3815 Chr7A. Critical recombinants spanning the *Pc94* locus delineated by PACE markers (red). Values indicate the number of recombinants observed between each marker. B. The physical region spans 3 Mbp with 5 predicted NLR genes (yellow; other genes in green). The *Pc94* region between markers P_As_26 and P_As_32 narrowed down through recombination with candidate genes and strongest candidate *G37624* showing exon structure and EMS mutagenesis locations. C. EMS mutants in *G37624* displaying susceptible infection types of 4 in response to *Pca* isolate 12SD80. D. Unrooted phylogenetic tree based on protein alignment of the zfBED domain of zfBED NLRs in clade 24 from diverse Poaceae species (Bailey et al. 2018) and *G37622* and *G37624* (labeled *AstNLR94*). The Poeae-specific, bootstrap supported clade is shown in light blue. E. Rooted phylogenetic tree based on protein alignment of the NB domain of zfBED NLRs in clade 24 from diverse Poaceae species (Bailey et al. 2018) and *G37622* and *G37624.* The bootstrap supported clade containing *AstNLR94* is indicated in grey, with the four Triticeae-specific and one Poeae-specific clades in blue, yellow, green, orange and light blue, respectively. (D and E) Red dots indicate supported branches at bootstrap values greater than 80% (1,000 bootstraps).

### Phylogenetic analysis

To perform phylogenetic analysis of N-terminal zfBED NLRs, we identified two sources of sequence information: (1) NLRs carrying N-terminal zfBED domains identified in the clade 24 MIC3 subclade from Bailey et al. (2018) (2) we interrogated Poeae genomic resources on NCBI and the *Avena* spp. pan-genome (Supplementary table 5) (Avni et al. 2025; Chen et al. 2025; Kamal et al. 2022; Maughan et al. 2019; NCBI Bioproject PRJNA1328942 2025; Pawłowicz et al. 2025; Wright et al. 2024; Zhang et al. 2025). For the latter, InterProScan (v5.23-62.0) using default parameters was used to identify protein domains. QKdomain scripts (https://github.com/matthewmoscou/QKdomain) were used to extract proteins with nucleotide binding domains with Pfam identifiers PF00027 and PF00931 and subsequently, proteins that contain zfBED domains (PF01163, PF01363, PF02892, PF04570, PF14634, PF00098, PF00642, PF00096, PS00028, PS00518, PS50089, PS50157, PS50178, PS50808, SSF57667, SSF57903, and SM00614). Manual curation of the identified zfBED NLRs was used to remove C-terminal zfBED fusions. Structure guided multiple sequence alignment was performed using MAFFT (v7.525) using parameters ‘--dash --maxinterate 1000 –globalpair’ (Katoh and Standley 2013). Model selection was carried out using modeltest-ng (v0.2.0) and identified models JTT+I+G4+F and JTT+I+G4 NB and zfBED domains, respectively, based on Bayesian Information Criterion (Darriba et al. 2020). Maximum likelihood phylogenetic analysis was performed using raxml-ng (v1.2.0) using parameters ‘--all –tree pars(100) --bs-trees 1000’ (Kozlov et al. 2019). RAxML input and analysis files are available on figshare (10.6084/m9.figshare.31100614). Phylogenetic trees were analyzed and visualized using iTOL (Letunic and Bork 2024).

### Cultivar genotyping and phenotyping

We PCR amplified and sequenced *G37624*, *G37621,* and *G37622* from *Pc94* cultivars as described above. Four seedlings of cultivars and non-carrier lines were phenotyped with genetically and pathogenically diverse isolates 90MN5B-1, 12NC29, 12SD80, 15FL1-2, 15MS7-1, 15MN23-1, 23FL26-1, 23SD76-1, 23MN60-1, and 23SD86-1 to compare phenotypic response of *Pc94* carriers and non-carriers. Pedigrees were assembled with the Helium website v2.1.0 (Shaw et al. 2014) using information from plant patents and the Pedigrees of Oat Lines (POOL) database (Tinker and Deyl 2005).

### Introgression size and location investigation

We performed a k-mer analysis to approximate *A. strigosa* introgression sizes in the *Pc94 A. sativa* cultivars and breeding lines. RNA from *A. sativa* and *A. strigosa Pc94*-containing accessions was extracted and sequenced as described above. Additionally, leaf RNAseq data from ‘Kanota’, ‘Sanfensan’, ‘Sang’, and ‘Victoria’ were downloaded from EMBL-EBI ENA (Kanota and Victoria Project: PRJNA422803, runs SRR6388184 and SRR6388185. Sang Project: PRJEB46365, runs ERR6323361 - ERR6323367. Sanfensan Project: PRJNA727473, runs SRR14934734 - SRR14934736) (Brabham et al. 2018; Kamal et al. 2022; Peng et al. 2022).

To identify regions that are potential introgressions, we used the K-mer Analysis Toolkit (KAT) v2.4.2 SECT tool (Mapleson et al. 2017) to determine k-mer coverage percentages of RNAseq data of each accession for each *A. strigosa* CIav 3815 gene coding sequence (n=37,787). We extracted genes with median k-mer coverage (%_non_zero_corrected values) greater than 89% in five *A. strigosa Pc94* carriers and maximum k-mer coverage less than 95% for *A. sativa Pc94* negative controls ‘Esker’, Kanota, Sang, Sanfensan, and Victoria. The filtered genes (n=9,411) were plotted based on their genomic location in CIav 3815. Regions with multiple genes with k-mer coverage near 100% were designated as introgressions and used to approximate introgression segment size. The KAT SECT output files and analysis R script are available on figshare (10.6084/m9.figshare.30896204).

We performed BLASTn searches for the *Pc94* candidate gene and adjacent zfBED NLRs in publicly available genome assemblies in GrainGenes: OT3098 v2, Sanfensan, and Sang, along with the *Pc94* carrier ‘Leggett’ with advance access granted by the pangenome consortium (Avni et al. 2025; Kamal et al. 2022; Peng et al. 2022; PepsiCo 2021; Yao et al. 2022). Follow up BLASTn analyses were run in Leggett and OT3098 v2 to determine the size and location of introgressions using all annotated genes in the predicted Leggett introgression region from the k-mer analysis and the adjacent region (Camacho et al. 2009). Chromosomal regions and contigs with high gene identity were aligned to the CIav 3815 assembly, visualized, and manually curated in Geneious Prime 2024.0.7 (https://www.geneious.com) using the Minimap2 v2.2.0 plugin (Li 2018). The contig order of the *Pc94* Leggett introgression region was reexamined with chromosome conformation capture (Hi-C) sequencing data generated from the PanOat project using Juicer v1.6 (Avni et al. 2025; Durand et al. 2016b) and visualized in R with strawr v0.0.92 (Durand and Shamim 2024), and the tidyverse (Wickham et al. 2019). The Leggett Juicer .hic output files for before and after rearrangement of contigs in the Pc94 introgression region (Figure 4B) are available on figshare (10.6084/m9.figshare.31058272).

### *Pc94* PACE diagnostic marker development

We designed the PACE markers using the assay design service by 3CR Biosciences (Essex, UK) (www.3crbio.com/free-assay-design). For the control gene, a primer pair was designed to amplify both *A. strigosa* EXO70A1 (AVATL.20003a.r1.6AG00000031060) and one homeolog of EXO70A1 on Chr4D (AVESA.00001b.r1.4Dg0002842) to make the assay functional in either species. Sequence alignment showed almost 100% conserved sequence within each species and homeolog group. A target gene primer pair was designed to amplify the *Pc94* candidate without off target hits in either species. PACE reactions were carried out with the same parameters described previously, except the assay mix consisted of 12 µM control gene forward primer, 30 µM control gene reverse primer, 12 µM target gene forward primer, and 30 µM target gene reverse primer in H_2_O. The raw PACE fluorescence data and R script for generating Figure 5 and Supplementary Figure 5 are available on figshare (10.6084/m9.figshare.31058098).

### QTL analysis

#### Genotyping by Sequencing

For QTL analysis, the two parents and core population of F_2_ individuals (F_2_-106 to F_2_-455) (n=314) were genotyped with genotyping by sequencing (GBS) by the University Minnesota Genomics Center. A double restriction digest (*PstI* and *MspI*) GBS protocol (Elshire et al. 2011) was performed with sequencing on AVITI (Element Biosciences) targeting 5-6 M 1 x 150 bp reads per sample. Reads were preprocessed with Trimmomatic as described above and aligned to the CIav 3815 genome assembly using bwa-mem2 (Vasimuddin et al. 2019). BCFtools mpileup and call were used to call SNPs with the multiallelic model. Sites with allele depth of <8 were recoded as missing, and loci filtered for CIav 3815 homozygous for the reference allele and CIav 1994 homozygous for non-reference allele. Ten samples were excluded for low read count (F_2__148, F_2__184, F_2__192, F_2__195, F_2__196, F_2__286, F_2__312, F_2__324, F_2__368, and F_2__395). Loci were quality filtered for heterozygotes <0.70, MAF>0.05, and missing genotypes <0.01. The dataset was pruned for sites with r2>0.9 in a 100kb window.

#### Linkage Map Construction

In R/qtl, markers were pruned based on segregation distortion using a P value threshold of 1e-10 and adjacent duplicate markers were dropped while ignoring missing data (exact.only=FALSE) (Broman et al. 2003). Linkage groups were assembled with max recombination fraction=0.15 and minimum LOD score=9. The original physical marker order was iteratively permutated in six marker sliding windows and reordered until the lowest crossing over values stabilized. Markers displaying many potential errors or double recombination events were removed after visual inspection of recombination graphs both before and after permutations. The linkage map and genotypes are available in Supplementary Table 6.

#### Phenotyping

F_2:3_ families were phenotyped with *Pca* isolates 12SD80 (two replications) and 90MN5B-1 (one replication) with 15-20 seedlings using the 0-4 IT scale as described above. Based on seed availability, 292, 233, and 221 F_2:3_ families were phenotyped for each isolate, respectively. The seedling ITs were converted to numerical values using the following scale: 0=0, ; (fleck) =0.25, ;n=0.75, 1=1, 1n=1.5, 2=2, 2n=2.5, 3=3, 3n=3.5, 4=4, and averaged to a single number for each F_2:3_ family for QTL analysis (Supplementary Table 7).

#### Composite Interval Mapping

We performed the QTL composite interval mapping (CIM) analysis with QTL Cartographer (Basten et al. 1994, 2004). For each trait, results from a five step forward stepwise regression with backward elimination were fed into CIM using model 6 using five background markers and 10 cM window size. A total of 1,000 permutations were performed with QKcartographer Python scripts to estimate α = 0.05 (https://github.com/matthewmoscou/QKcartographer). Logarithm of odds (LOD) test statistics for H0:H3 (testing for additive and dominance effects) were processed and graphed with QKcartographer and ggplot2 in R.

## Results

### QTL analysis of *Pc94*

At the beginning of the project, we verified the placement of the *Pc94* locus in previously constructed linkage maps and located the associated markers on the *A. strigosa* CIav 3815 genome assembly to determine the region targeted for recombination screening. In the GBS CIav 3815 x CIav 1994 RIL population map (Latta et al. 2019), the *Pc94* locus from Kremer et al. (2001) was binned with distal markers Oat18480.1, Oat38313.1, Oat408.1, Oat5006.1, and Oat18466.1 on LG7 at position 136.1 cM. To verify the putative location of *Pc94*, we performed a preliminary QTL analysis using the 2019 linkage map and original RIL phenotypic data (Wise et al. 1996), which confirmed the placement of a major locus at position 136.1 (LOD 16-23.5) (Supplementary Fig. 2). The markers spanned an 800 kb region between 597,079,200 bp and 597,957,290 bp on Chr7A in *A. strigosa* CIav 3815, adjacent to the end of the assembled chromosome. During manual curation of the region, we combined two separately annotated genes into a single gene (*G37625/G37626*) as the two halves made up one full NLR and were equally highly expressed in unchallenged leaf tissue. The region contains five annotated NLR genes, all of which have integrated zfBED domains. We designed PACE markers to screen the region for recombinants.

### *Pc94* fine mapping

To refine the *Pc94* region and identify candidate genes, we screened 851 F_2_ plants with PACE markers to detect recombination events, yielding 47 recombinants. F_2:3_ family phenotypic results for the recombinant lines based on three different *Pca* isolates narrowed down the region to 500 kb between markers P_As_26 and P_As_32, eliminating two of the five candidate NLRs previously identified in the region (Fig. 1A, Supplementary Table 8). RNAseq of two homozygous recombinants between markers P_As_07 and P_As_32 eliminated candidate NLR *G37625/G37626*, as resistant F_3_-233 did not express *G37625/G37626* and susceptible F_3_-474 expressed *G37625/G37626* (Supplementary Fig. 3A). Although *G37624* showed markedly lower expression (Supplementary Fig. 3B), it was PCR amplified and sequenced from resistant F_3_-233 and not from susceptible F_3_-474, demonstrating recombination between the two genes in both recombinant accessions.

This narrowed the *Pc94* region to five candidate genes, including two NLRs (*G37622* and *G37624*) and one Cystine-Rich Receptor-Like Kinase annotated as a homolog of Arabidopsis CRK10 (*G37621*). We focused on the CRK and two NLRs as the main candidates as they were only present in the resistant parent’s assembly and these gene families are associated with disease resistance.

We screened an additional 860 F_2_ plants and 340 F_4_ plants specifically for recombinants in the refined region, identifying zero recombinants between P_As_10 and P_As_07 and two F_2_ recombinants between P_As_26 and P_As_10. One of these recombinants (1239) was successfully advanced to homozygosity in the F_3_ generation. The CRK10 homolog (*G37621*) was PCR amplified from susceptible homozygous recombinant F_3_-1239; therefore it was ruled out as a candidate for *Pc94* under a single gene model. Based on the recombination screen, we narrowed the *Pc94* locus down to two candidate genes, NLRs *G37622* and *G37624*.

### EMS mutagenesis screen

We used a mutagenesis approach to determine if any of the candidate genes in the *Pc94* genetic region were required for *Pc94*-mediated resistance. An EMS mutagenesis screen yielded 17 susceptible *M*_2_ plants from eight individual *M*_1_ panicles. Two distinct mutations were observed in NLR *G37624*, with four lines (EMS24-093-1, EMS24-340-1, EMS24-340-2, and EMS24-340-3) exhibiting g.C784T and one line, EMS24-004-1, exhibiting g.G2251A (Fig. 1B). These point mutations lead to amino acid substitutions in the predicted zfBED domain (p.T197I) and the predicted disease resistance protein winged helix domain (pfam PF23559) (p.G646S). The zfBED domain mutants did not have mutations in NLR *G37622* or *CRK10* homolog *G37621*. However, EMS24-004-1 also exhibited a missense mutation in *G37622* in the LRR domain (g.C3888T, p.L1296F) and a missense mutation in *G37621* (g.C1061T, p.A354V). The two observed amino acid substitutions in *G37624* alter amino acid classes — one from polar uncharged to hydrophobic and the other from glycine (special-case residue) to polar uncharged — whereas substitutions in *G37622* and *G37621* in EMS24-004-1 involve exchanges between hydrophobic amino acids.

For the remaining susceptible *M*_2_ mutants, one mutant, EMS24-234-1, did not have homozygous mutations in any of the three genes and its susceptibility was likely the result of mutations affecting other genes involved in *Pc94*-mediated resistance. Susceptible plants derived from four *M*_1_ panicles did not contain any of the three candidate genes and were determined to be derived from rare off-types in the original *Pc94* differential seed lot. We observed rare susceptible plants in the phenotypic screen’s unmutated *A. sativa Pc94* differential control seedlings which when tested also did not contain the three candidate genes.

To understand if natural variation in *Pca* affected the phenotype of *Pc94* loss-of-function mutants, we tested homozygous *M*_3_*G37624* mutants with nine diverse *Pc94* avirulent isolates. In most cases, all *G37624* mutants displayed susceptible ITs of 3 or 4 that matched the susceptible control showing complete susceptibility (Fig. 1C, Supplementary Table 9). For two of the nine isolates, EMS24-004-1 exhibited a slightly less susceptible IT than the other mutants (isolate 90MN5B-1: 2 vs 4 and isolate 12NC29: 3 vs 4) (Supplementary Fig. 4), suggesting the mutation in the predicted disease resistance protein winged helix domain is not as disruptive to *Pc94* resistance as the zfBED domain mutation.

The critical recombinants and mutagenesis screen demonstrate *G37624* is necessary for *Pc94* resistance, however sufficiency has not yet been determined, therefore we designated *G37624* as *AstNLR94*, *Avena strigosa NLR required for Pc94 resistance*.

To understand the origin and relationship of *AstNLR94* and *G37622* to previously cloned resistance genes encoding zfBED NLRs, maximum likelihood phylogenetic analysis was performed with Poaceae zfBED NLRs. In analyzing the NB domains, *AstNLR94* and *G37622* were found to belong to a clade of zfBED NLRs that includes a Poeae-specific and four Triticeae-specific clades, indicating substantial expansion of this clade in Triticeae species (Fig. 1E). One of the Triticeae-specific clades includes the cloned wheat powdery mildew resistance gene *Pm6Sl* (Ma et al. 2024b), indicating a common ancestor for these zfBED NLRs. In contrast, analysis of the smaller zfBED domain found a single supported clade including only Poeae species that corresponds with the NB clade including Poeae species (Fig. 1D).

### Confirming *Pc94* candidates in *A. sativa* germplasm

Next, we PCR amplified and sequenced the *Pc94* candidate genes in *A. sativa Pc94* carriers. *AstNLR94*, *G37621*, and *G37622* were confirmed in the following cultivars: ‘Leggett’, ‘ND Carson’ ‘MN Pearl’, ‘Rushmore’, ‘Saddle’, ‘SD Buffalo’, ‘Sumo’, and ‘Warrior’, consistent with *Pc94* carrier phenotypic results from seedling tests (Supplementary Table 10) and pedigree information (Fig. 2) (Caffe-Treml 2021a, b, c, d, 2023; McMullen 2025; Mitchell Fetch et al. 2007; Smith 2020). None of the genes amplified in AAC Douglas despite containing Leggett in its pedigree three generations prior (Mitchell Fetch et al. 2023).

**Fig. 2.**
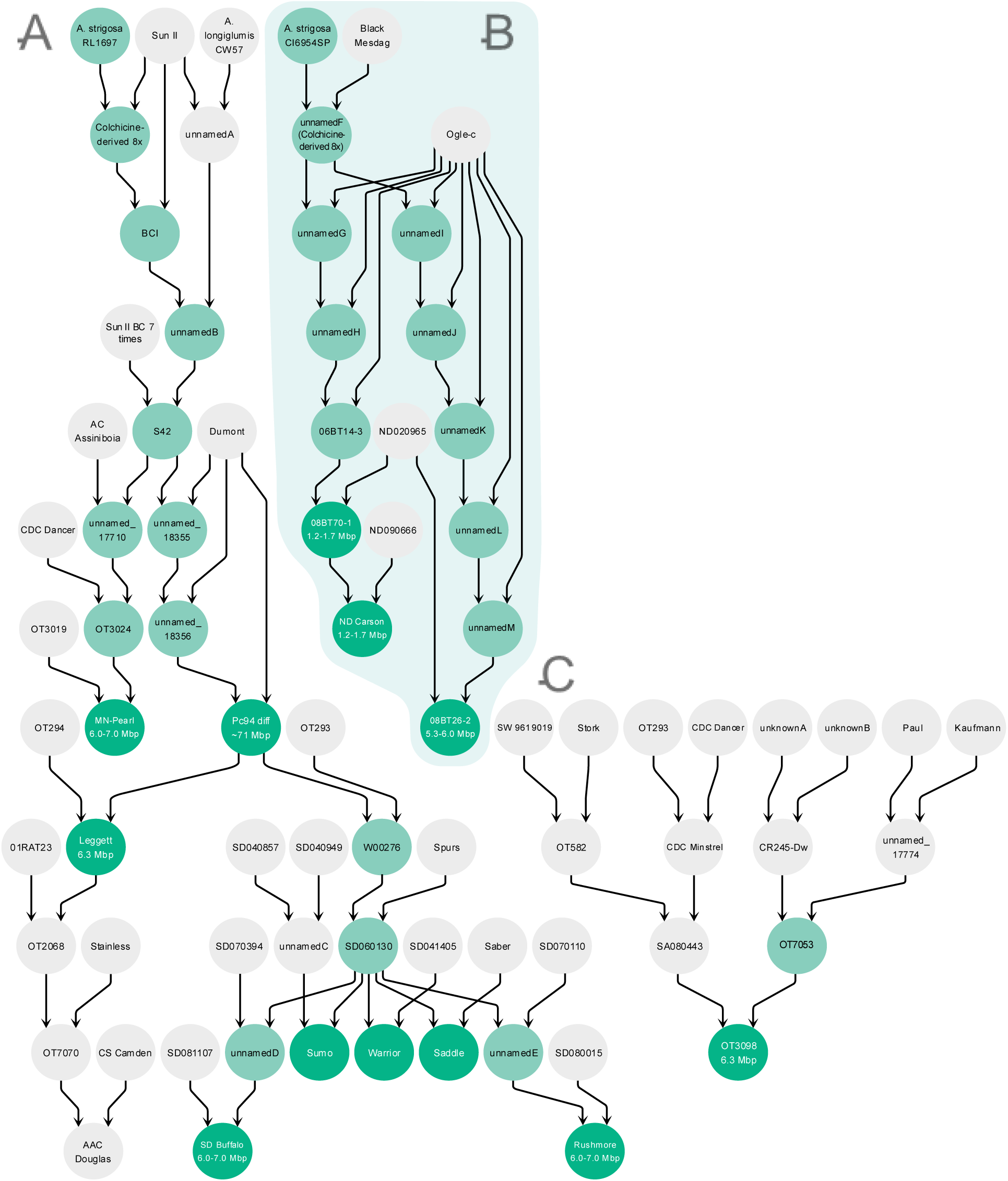
Pedigrees of the *Pc94* introgressions. A. 1^st^ *Pc94* introgression B. 2^nd^ *Pc94* introgression C. OT3098 pedigree. Accessions in dark teal are confirmed *Pc94* carriers via PCR and phenotyping, accessions in light teal are hypothesized as *Pc94* carriers based on pedigree or phenotypic information. The introgression size estimates based on RNAseq or genome assembly analysis are included under accessions. Note the seven backcrosses of Sun II to produce S42 were condensed in A for a more compact figure. Names are consistent with the POOL oat pedigree database where applicable and unnamed accessions from patent certificates and the scientific literature were assigned distinguishing letters.

### Introgression of *Pc94* region in *A. sativa*

To understand if *A. strigosa* introgression region sizes varied in different *Pc94* introgression lineages and cultivars, we estimated the approximate size of the introgressions for many of the *Pc94* carriers using RNAseq data. We determined k-mer coverage of *A. strigosa* genes in leaf RNAseq data from *Pc94* carriers and non-carriers and plotted the k-mer coverage of genes that had greater than 89% median k-mer coverage in *A. strigosa* accessions with *Pc94*, but less than 95% k-mer coverage in any non *Pc94*-carrier *A. sativa*s, searching for regions in each *A. sativa Pc94* carrier accession with nearly 100% k-mer coverage of genes from *A. strigosa* (Figure 3). All the *Pc94* carriers contained introgressions from *A. strigosa* Chr7A without additional chromosomal regions. The largest introgression, ∼71 Mbp, was found in the *Pc94* differential line. The introgression size estimate of the cultivars tested from the first introgression event were the same, with 6.0-7.0 Mbp introgressions in MN Pearl, Leggett, Rushmore, and SD Buffalo. The introgression in 08BT26-2 was estimated to be slightly smaller, at 5.3-6.0 Mbp. The smallest introgressions were in 08BT70-1 and its progeny ND Carson, at 1.2-1.7 Mbp, with *AstNLR94* as one of the first genes detected at the introgression boundary. The introgression estimation gives ranges because only a fourth of the genes passed filtering to be used in the analysis and in some cases the regions between informative genes were large due to repetitive elements. Using this analysis, we estimated the *Pc94* carriers’ introgressions range in size from 1.7 Mb to 71 Mb of *A. strigosa* Chr7A DNA.

**Fig. 3.**
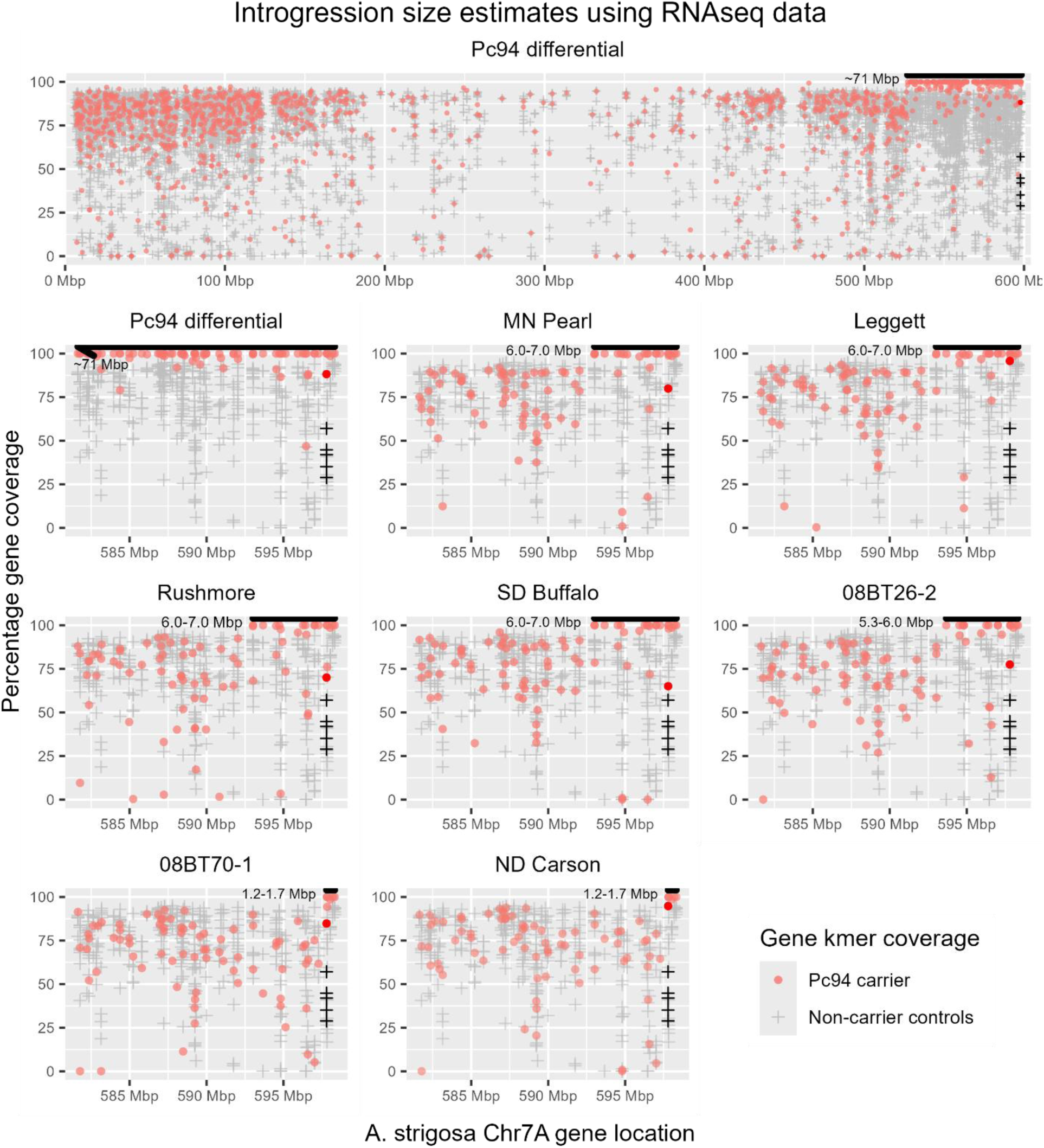
*Avena strigosa* introgression size estimates using RNAseq data and k-mer analysis for 8 *Avena sativa Pc94* carriers. Estimated introgression sizes are shown with a thick black line at the top of each plot. For each gene in *A. strigosa*, red circles are the k-mer coverage of *Pc94* carriers, and gray plus marks are the k-mer coverage of five non *Pc94* carrier controls. *AstNLR94* is indicated by the darker red and black symbols.

To determine the location and exact size of the *Pc94* introgression in the Leggett genome, we performed BLAST analyses with the genes from the RNAseq predicted introgression region in CIav 3815 on the Leggett genome assembly. We discovered a 6.3 Mbp introgression region split between the beginning of Chr7D, the end of Chr2A, and three small unplaced contigs (Figure 4A). Reexamination of the HiC data for these regions showed a small but likely misassembly, with strong evidence for a contiguous sequence at the end of Chr2A that matches the CIav 3815 Ch7A assembly (Figure 4B). A contiguous introgression is more logical because the introgression size is conserved in all cultivars tested from the first introgression. If it was split on two separate chromosomes, the segment without *Pc94* would likely have been segregated out in breeding due to random segregation if not by selection.

**Fig. 4.**
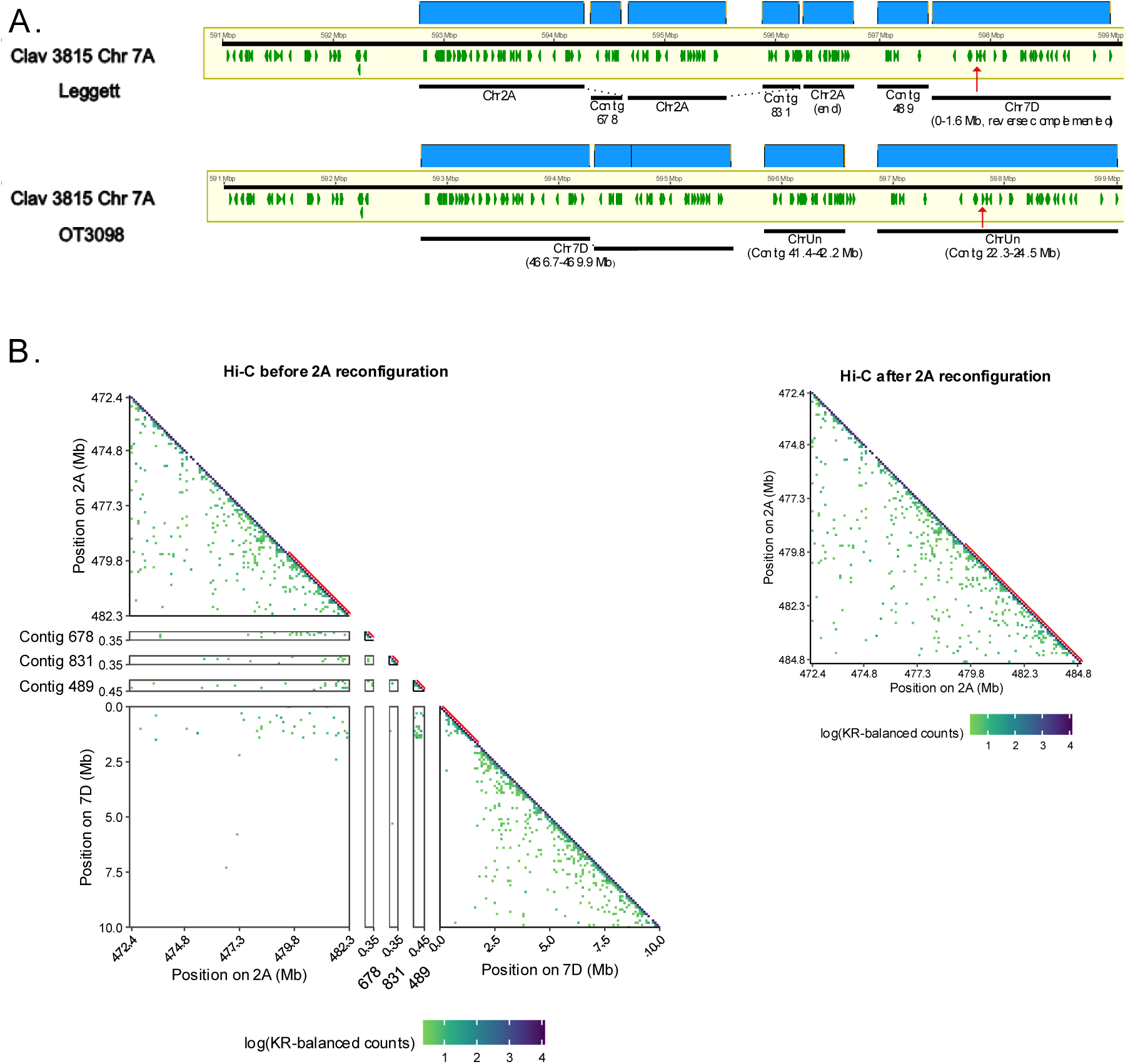
Evidence for contiguous *Pc94* introgression region in *Avena sativa*. A. Layout of contigs from *Pc94* carriers Leggett and OT3098 with nearly 100% identity to *A. strigosa* CIav 3815 Chr 7A *Pc94* region. The red arrow designates *AstNLR94*. B. Chromatin contact maps from Leggett HiC data before and after reconfiguration of Chr2A to add contigs 678, 831, and 489, as well as a 1.6 Mb contig originally scaffolded on the beginning of Chr7D. The red line designates introgression DNA from *A. strigosa*.

Unexpectedly, we also discovered *Pc94* in the PepsiCo OT3098v2 genome. The OT3098v2 assembly contains the same *A. strigosa* introgression size as Leggett (Figure 4A). In this assembly, the *A. strigosa* introgression is split between two contigs from the concatenated contig sequence Chr00 (ChrUn) and a section of Chr7D from 41.1-42.2Mbp. The introgression regions of both Leggett and OT3098 share almost 100% identity with CIav 3815, with most differences attributable to possible sequencing errors. When we investigated OT3098 further, its phenotypic response was consistent with other *Pc94* carriers (Supplementary Table 11).

As part of his introgression efforts with *Pc94* at the USDA Cereal Disease Laboratory, Howard Rines investigated the relative genomic locations of the *Pc94* introgressions (previously unpublished data). In addition to 08BT26-2 and 08BT70-1, he originally obtained five other independently derived *Pc94* lines from backcrossing the *A. strigosa* CI6954SP x Black Mesdag colchicine-derived octoploid to Ogle-c (Cornell reselection of Ogle). Five of the seven independently derived lines when crossed with either Leggett or the *Pc94* differential line yielded no susceptible F_2_ plants as would be expected if the genes were in the same genomic location. Crossing 06BT14-3 (parent of 08BT70-1, ND Carson lineage) with Leggett and the differential *Pc94* line resulted in segregating F_2_ populations of 83R:5S and 151R:6S, respectively. These data fit a 15:1 ratio expected for two independent *R* genes (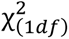= 0.05, *P* = 0.82 and 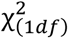 = 1.57, *P* = 0.21), demonstrating two separate *Pc94* genomic locations from the introgressions used in developing modern cultivars. The location of the *Pc94* introgression in the ND Carson lineage is yet to be determined. It is notable that in 6 of the 8 introgressions (1/1 Canadian and 5/7 USDA), the introgression incorporated onto *A. sativa* Chr2A. Homology between the two regions may be directing the introgression to Chr2A.

### *Pc94* Marker Development

We designed an improved, high-throughput *Pc94* marker for use by breeders and geneticists. We developed a presence/absence PACE genotyping assay in *AstNLR94* which positively identified all 12 *A. sativa Pc94* carriers from both introgressions. There was no off-target amplification in 84 diverse *A. sativa* non-carrier cultivars and lines (Supplementary Table 11). Additionally, the marker amplified all eight *A. strigosa AstNLR94* carriers in a 93-accession diversity panel without off target amplification as verified by PCR amplification and sequencing (Supplementary Fig. 5). In this assay, the control amplifies a conserved region of EXO70A1 in *A. strigosa* or one homeolog of EXO70A1 on Chr4D in *A. sativa*, enabling use of the assay in either species. Amplification for the control gene (FAM signal) signifies PCR was successful but *Pc94* is absent, and amplification of both alleles (FAM and HEX signals) indicates *Pc94* is present (Figure 5). To understand why the previous Single Base Extention (SBE) (Chen et al. 2007; Chong et al. 2004) marker was considered uninformative, we examined its genomic location and found it lies within the gene AVATL.20003a.r1.7AG00000037209 at 582 Mbp Chr7A in the CIav 3815 assembly, 16 Mbp away from *AstNLR94*. The SBE marker was developed using S42, one of the oldest accessions in the first *Pc94* introgression lineage. However, the SBE marker has limited utility because it fails to detect accessions carrying smaller introgressions, such as Leggett, OT3098, and other modern *Pc94* cultivars. This highlights the importance of developing a diagnostic marker within the causal gene.

**Fig. 5.**
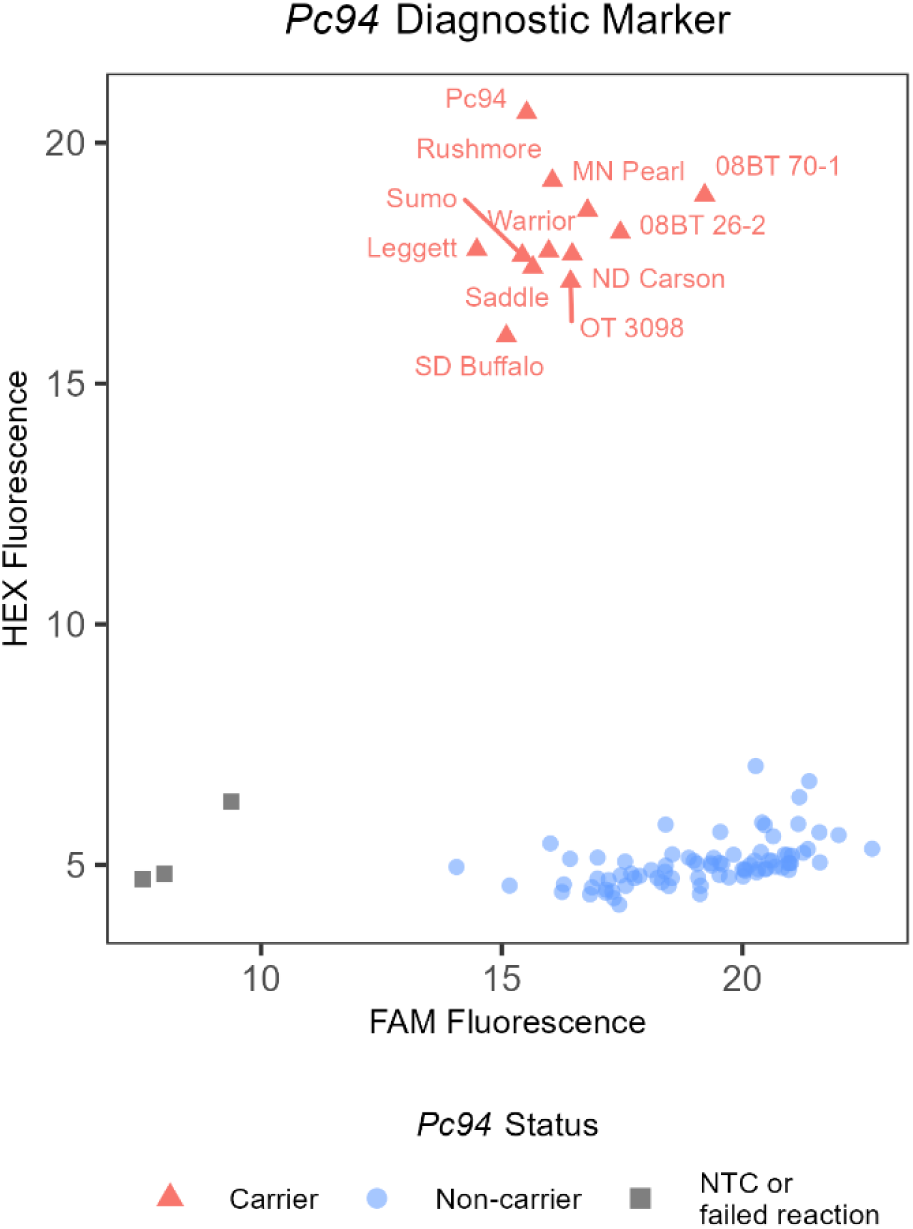
*Pc94* PACE fluorescence cluster plot. *Avena sativa Pc94* carriers are labeled as red triangles.

### QTL analysis identifies a minor locus in CIav 3815 background

During the evaluation of F_2:3_ families, variation in the degree of resistance and susceptibility was observed several times and warranted further investigation into other potentially important resistance loci segregating in the population. For example, we observed resistant families with ITs from ; (fleck) to 2n and susceptible families from 3 to 4. Furthermore, several recently collected USDA *Pca* survey isolates completely virulent to the *Pc94* differential line (IT of 4) demonstrated less severe ITs of 2 and 3 on CIav 3815 (Supplementary Table 12) suggesting additional resistance in this accession.

To conduct a QTL analysis for minor *R* loci in the parents, we first constructed a linkage map for the core F_2_ population (n=304), resulting in seven linkage maps with 566 GBS-derived markers ranging in size from 48-116 markers and 123-184 cM (Supplementary Table 6). Comparisons of markers on the linkage map with the CIav 3815 genome assembly show good marker order agreement and large centromeric zones of reduced recombination (Supplementary Fig. 6). In the QTL analysis using F_2:3_ family IT averages, one locus on Chr4A (5-12 cM) was significantly associated with resistance to isolate 90MN5B-1 and two replications of 12SD80 with LOD scores of 5.1-7.2 explaining 2.8-4.1% of the phenotypic variation (Figure 6, Supplementary Fig. 7). The associated markers are located on Chr4A from 19.6-33.7 Mb. Additional QTL were unique to the different inoculations and may represent additional isolate specific loci, for example for Chr7A with associated marker at 351.9 Mb for 90MN5B-1 (Table 1). As expected, the *Pc94* locus at the end of Chr7A was the major peak for all three traits with LOD scores of 62.2-95.7 explaining 54-82% of the variance.

**Fig. 6.**
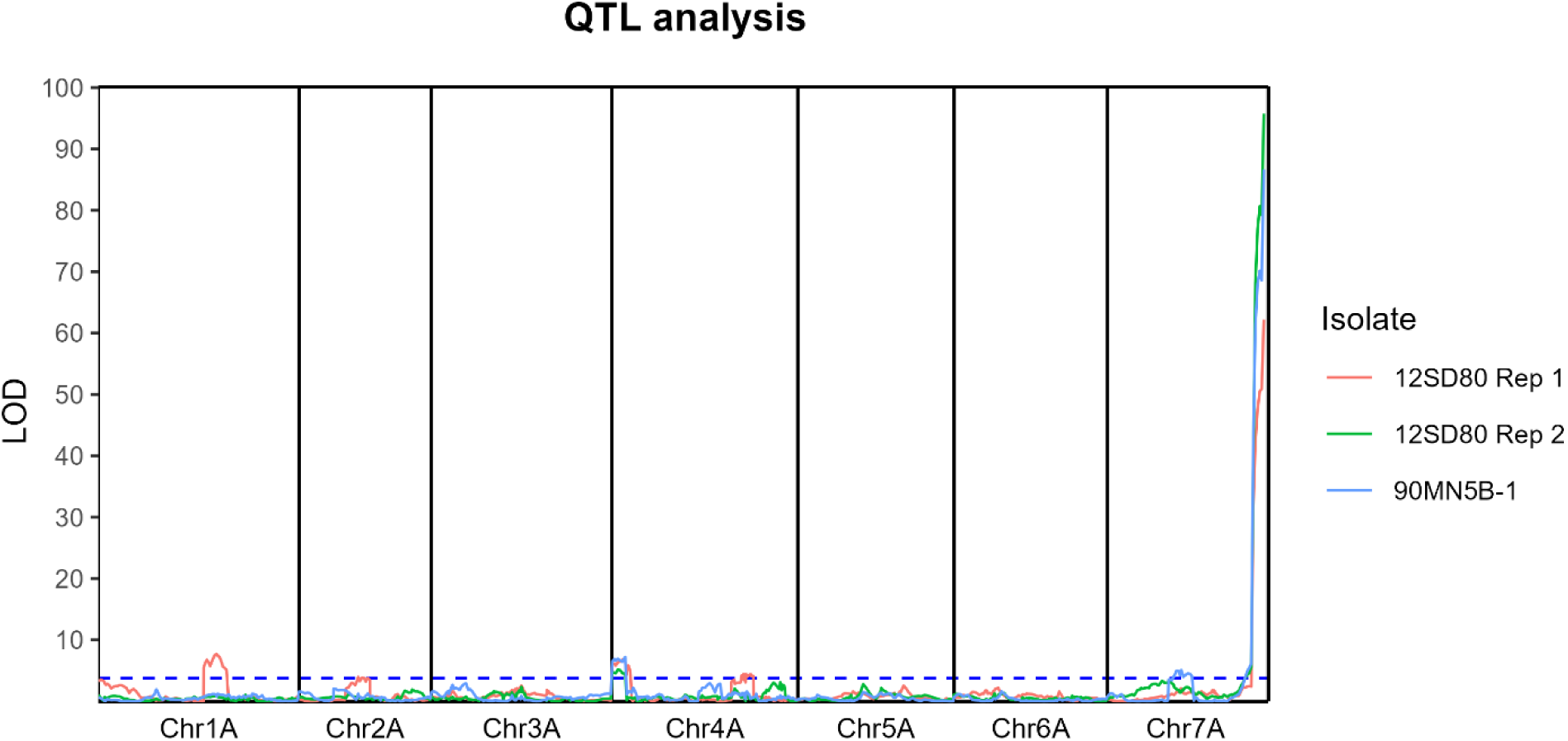
Logarithm of Odds (LOD) plot from QTL analysis of F_2:3_ families segregating for *Pc94*. The dotted blue line is the significance threshold at α = 0.05 for all three isolates.

**Table 1.**
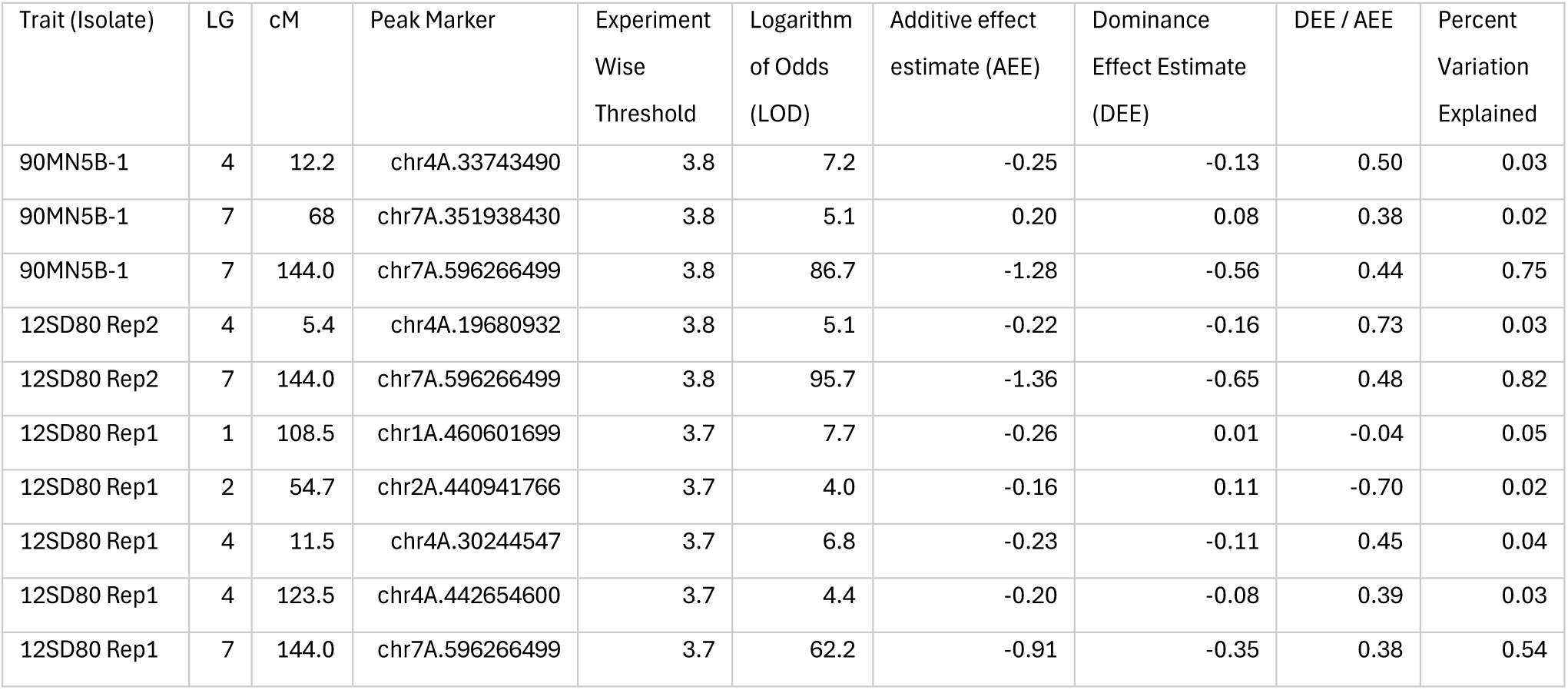
Significant QTL from QTL analysis of F_2:3_ families segregating for *Pc94*. Markers are named by their position in the CIav 3815 genome assembly.

## Discussion

*Pc94* is the first *R* gene identified to a single candidate gene in oat, a major step forward for resistance breeding in this important cereal crop. The work was facilitated by the ease of fine mapping and genome assembly in the cultivated diploid *A. strigosa* donor. Fine mapping efforts in *A. sativa* are complicated by the complex hexaploid genome with abundant structural variation and homologous regions (Avni et al. 2025; Zhang et al. 2025). Furthermore, when *Pc* genes derived from other oat species are mapped in *A. sativa* populations, the introgression regions may also exhibit suppressed recombination (Bekele et al. 2025; Wight et al. 2024). Our work provides several independent lines of evidence that *AstNLR94* is necessary for *Pc94*-mediated resistance including recombination mapping, mutagenesis, and gene expression analysis. Sufficiency of *AstNLR94* is still to be demonstrated before it can be designated as *Pc94*.

A major advance from this work was the development of a *Pc94* PACE diagnostic marker, which fits into the *Pc* gene screening workflow of modern genotyping laboratories and is suitable for MAS. Designing a specific marker based on a small region within an NLR gene was challenging, taking many iterations due to off-target amplification in a very large and variable gene family. The final diagnostic marker was designed in the highly variable LRR domain. The *Pc94* marker is presence/absence: a limitation because it will not distinguish between *Pc94* homozygotes and heterozygotes. Differentiation between zygosities is useful for breeders and scientists to determine if *Pc94* is fixed in lines without having to phenotype or genotype the next generation for segregation of resistance. However, a standard PACE approach was not feasible because there was no suitable alternate allele for designing a competitive forward primer; the *Pc94* locus is either present or absent and depending on the introgression origin, may be in a different chromosomal location. Nevertheless, this *Pc94* marker is considerably improved over previous markers and should not exhibit recombination with *Pc94* resistance.

The *Pc94* locus has a convoluted naming and mapping history since its description in the 1950s. An initial source of confusion seems to have arisen from *Pc94*’s presence in several distinct *A. strigosa* accessions and additional minor resistance genes in these backgrounds. The accession CIav 3815 was first described to carry *Pc19* (Simons et al. 1959) and a secondary, unlinked locus *Pc30* (Marshall and Myers 1961; Simons et al. 1978). Sadanaga and Simons report a stable introgression of one of these loci into *A. sativa* (1967), however, there is no record of the introgression being used in cultivar development. In the 1990’s, CIav 3815’s resistance was mapped as the “*Pca*” locus (Rayapati et al. 1994) then described as five tightly linked isolate specificities designated *Pc81-85* (Wise et al. 1996). Depending on the *Pca* isolate used, the resistance was also noted to be influenced by other dominant loci. *Pc81-85* names were assigned without investigation of their relation to *Pc19* and *Pc30* or potential precedence in naming (Park et al. 2022; Yu and Wise 2000). Concurrently, an *R* locus from *A. strigosa* RL1697 (designated as *Pc94*) was introgressed into *A. sativa* (Aung et al. 1996) and released in the Canadian cultivar ‘Leggett’ (Mitchell Fetch et al. 2007). Inspired by this introgression, USDA scientists transferred a promising *R* locus from *A. strigosa* CI6954SP into *A. sativa*. However, they eventually concluded this locus was likely the same as *Pc94* found in RL1697 based on several lines of evidence, including marker linkage, mutual suppression by *Pc38*, and shared susceptibility to a new virulent rust isolate (Rines et al. 2007).

Allelism tests for the genetic resistance in CIav 3815, RL1697, and CI6954SP showed no segregation of resistance in over 800 RL1697 x CIav 3815 F_2_ seedlings or 100 CI6954SP x CIav 3815 F_2_ seedlings (Park et al. 2022; Rines et al. 2007). Despite the *Pc19* designation having precedent, this study refers to the locus as *Pc94* for consistency with the longstanding name used in the oat breeding and pathology community.

Results from this study support the original work with CIav 3815 suggesting one major gene (*Pc94*) with a minor secondary locus (*Pc30*), which we placed on the beginning of Chr4A. *Pc30* was not transferred into *A. sativa*. Interestingly, there is a nonsignificant peak in the vicinity of *Pc30* in the preliminary QTL analysis using the original 100 individual RIL population despite the small size and binary phenotyping data. We were unable to repeat the results from the 1990’s showing five closely linked, differentiating loci (*Pc81-85*) in the RIL population in preliminary tests with several isolates nor did we detect evidence of this in the larger F_2_ population. When the RIL population was genotyped with the recombination PACE markers, no recombination was observed in the NLR region delineated by the significant markers from the original QTL analysis, suggesting that the differences in specificities were likely due to other segregating loci in the background of the population in addition to the inherent subjective nature of phenotyping at the boundary of resistant/susceptible classification (2 vs 3 infection type) where some of phenotypes lie. However, considerable time had passed since the initial work on the RIL population that may have impacted purity and identity of several RILs in our possession, prompting us to develop new populations.

*Pc94* resistance appears to have resisted breakdown for a relatively long time, about 15 years. However, this may have been due to relatively limited acreage share of *Pc94* cultivars and not due to special properties of *Pc94*. The Canadian cultivar Leggett, released in 2007, stacked *Pc94* and *Pc68;* however, virulence to *Pc68* had increased dramatically in the eastern prairie region of Canada by the early 2000s (Chong et al. 2008), effectively placing most of the selection pressure on *Pc94*. Insured Leggett acreage peaked in the prairie regions of Canada in 2010 at 111K acres (11% cultivar share) which then steadily decreased to less than 1% in 2017 (Canadian Grain Commission 2024). No other widely grown variety in that region contains *Pc94* as far as we could determine. We note that AAC Douglas contains *Pc94* in its pedigree, however it tested negative for *AstNLR94* and other candidate genes in the *Pc94* region via PCR amplification and likely contains resistance from other sources (e.g. *Pc91*, ‘Stainless’) (Mitchell Fetch et al. 2023). The unexpected discovery of *Pc94* in OT3098 highlights the possibility it is unknowingly in other Canadian breeding material and cultivars, however cultivars ‘AAC Almonte’, ‘AAC Justice’, ‘AAC Nicolas’, ‘AAC Oaklin’, ‘AAC Roskens’, ‘AC Assiniboia’, ‘Cascade’, ‘CDC Endure’ ‘CS Camden’, ‘Exeter’, ‘Garry’, and ‘Optimum’ all tested negative for the *Pc94* marker. Virulence to *Pc94* spiked in the Northern US in 2022, soon after the release of *Pc94* in several cultivars from multiple breeding programs. Increased levels of virulence to *Pc94* in Canada since 2020 may be due to wind-dispersed inoculum from the Northern US or from the presence of *Pc94* in another widely grown but unsampled Canadian cultivar.

The new genome assemblies along with the interesting history of several independent introgressions of *Pc94* offer a unique opportunity to trace introgression size through generations and investigate introgression location in independent lineages. Cultivar development has decreased the first introgression from ∼71 Mbp in the *Pc94* differential to 6.0-7.0 Mbp. Interestingly, all cultivars included in the RNAseq analysis from the first introgression have the same 6.0-7.0 Mbp signal, even though there are varying numbers of generations separating each from the original introgression and MN Pearl was derived from an earlier line in the first introgression pedigree (S42) than the *Pc94* differential (Figure 2). Recombination suppression may be preventing further shortening of the introgressed region, which contains ∼150 predicted *A. strigosa* genes. The introgression in the USDA germplasm line 08BT26-2 is slightly smaller at ∼5.3-6.0 Mbp but no cultivars have been derived from it. These introgressions are situated at the end of Chr2A. The other tested USDA *Pc94* introgression line, 08BT70-1, and its progeny ND Carson have the smallest introgression (∼1.2-1.7 Mbp, ∼25 genes), which is located at an undetermined genomic location based on independent segregation. Other cases of independently characterized and introgressed, yet redundant, wild relative-derived *Pc* genes may occur and be complicated by distinct introgression locations in *A. sativa*, further confusing efforts to resolve *Pc* gene nomenclature and reduce redundancy.

*Pc94* can be clearly traced through pedigrees due to its recent introgressions into *A. sativa*. It was only discovered in one unexpected accession, OT3098, a breeding line from the Crop Development Centre (CDC) at the University of Saskatchewan. OT3098 was used for the first *A. sativa* genome assembly (Figure 2). There is strong evidence *Pc94* in OT3098 was derived from the first introgression. The introgression size in the OT3098 v2 assembly exactly matches that of Leggett and is larger than the introgressions in 08BT26-2 and 08BT70-1. The breeding cross for OT3098 (OT7053 x SA080443) was made in 2009, two years before Howard Rines began sharing his *Pc94* introgression with breeders and the CDC never used Rines’ *Pc94* breeding lines in the program (Aaron Beattie, personal communication). However, OT3098’s introgression is split between several different contigs in its genome assembly, none of which is located on Chr2A. Due to its source from the first introgression, this is likely a misassembly with the true location at the end of Chr2A, but further investigation is needed to clarify exactly what has occurred (unfortunately lack of access to the Hi-C data limited our ability to resolve the scaffold in the OT3098 assembly). Large repetitive regions seem to make this introgression region especially complicated to assemble. As to the parental source of *Pc94* in OT3098: Leggett, OT3098, and the Agriculture and Agri-Food Canada-sourced OT7053 parent shared the same OCR resistance/susceptibility pattern: resistant to most tested isolates, but susceptible to isolate BRBG-94 (Supplementary Table 13) (data curtesy of Aaron Beattie and Kirby Nilsen). Therefore, OT7053 is likely the immediate source of *Pc94* in OT3098. There is not enough data to trace *Pc94* further up the pedigree, but CR245-Dw is a possibility as its parentage is unknown.

*AstNLR94* is a member of the zfBED NLR family, a widespread and monophyletic NLR class in monocots. An analysis found zfBED NLR encoding genes in 7 of 9 grass genomes, with the exception of maize and sorghum (Bailey et al. 2018). Cloned zfBed NLR-IDs confer resistance to barley leaf rust (*Rph15*), wheat powdery mildew (*Pm6Sl*), wheat stripe rust (*Yr5/YRSP* and *Yr7*), and bacterial blight and bacterial leaf streak in rice (*Xa1* and *Xo1*) (Chen et al. 2021; Ma et al. 2024a; Marchal et al. 2018; Read et al. 2020a; Read et al. 2020b; Yoshimura et al. 1998). Interestingly, unlike other types of NLR-IDs characterized to date, these genes have not been shown to require a paired executive NLR to trigger an immune response (Ma et al. 2024b; Marchal et al. 2022). Therefore, we hypothesize *AstNLR94* will prove sufficient for *Pc94* resistance.

The mechanism by which zfBED IDs contribute to NLR-mediated resistance remains unknown. One hypothesis proposes zfBED IDs act as decoys for pathogen effectors that would normally target host transcription factors containing zfBED domains for DNA binding (Grund et al. 2019), however, direct effector-zfBED domain interaction has not yet been demonstrated. The XA1 zfBED domain forms complexes with bacterial transcription activator-like (TAL) effectors through a transcription factor intermediary (Yoshihisa et al. 2022). However, XA1 zfBED domain is identical to that of XO1, yet *Xo1* confers resistance to an additional pathovar of *Xanthomonas oryzae* (Read et al. 2020b). In wheat, the YR5/YRSP alleles are almost identical; YRSP has a truncated LRR domain compared to YR5 and protects against considerably fewer stripe rust races (Marchal et al. 2018). In both instances, recognition specificity is not solely due to the zfBED domain, although the zfBED domains are necessary for resistance. Likewise, an amino acid substitution in the ASTNLR94 zfBED domain was sufficient to induce susceptibility for all isolates tested, while the second mutation in a different domain showed some isolate specificity. The identification of *AstNLR94* offers an additional gene for elucidating resistance mechanisms in the zfBED NLR family to diverse diseases, which may help inform *R* gene engineering efforts.

Determining the corresponding *Pc94* pathogen effector, *AvrPc94*, will be a focus of future work. The sexual *Pca* population in North America lends itself well to effector genome wide association studies, and previous work has pinpointed candidate effectors for other *Pc* genes (Hewitt et al. 2023; Miller et al. 2021). Adding *Pc94* virulent isolates to the original datasets would enable identification of *AvrPc94* candidates. Testing protein-protein interactions between *AstNLR94* and top *AvrPc94* candidates using AlphaFold-Multimer could serve as a complimentary analysis and provide insight into the physical interaction domains of zfBED NLRs and their corresponding effectors. Another interesting avenue of inquiry is elucidating the mechanism of *Pc94* suppression by *Pc38* as perhaps the Pc94 zfBED domain is affected in the interaction with Pc38. As *R* gene engineering becomes more feasible, the interactions between *R* genes will need to be better understood and taken into consideration for successful deployment of novel resistances from wild relatives or gene editing.

This study presents the most in-depth characterization of a resistance gene candidate and interspecific introgression in oat to date. The newly developed *Pc94* marker will greatly facilitate tracking the locus in germplasm and cultivars as well as marker assisted selection for breeders. Careful deployment of *Pc94* by stacking with other qualitative and quantitative resistance should help prolong its effectiveness in oat growing regions where it is not currently in use.

## Statements and Declarations

### Author contributions

Conceptualization: EPM, MM, and SK; Methodology: EPM, PJM, MM; Formal analysis and investigation: EPM, RS, PJM, MM; Visualization: EPM and MM; Writing - original draft preparation: EPM; Writing - review and editing: EPM, PJM, RS, ENJ, MM, SK; Funding acquisition: MM and SK; Project Administration: EPM; Supervision: MM and SK.

## Acknowledgements

We gratefully acknowledge the late Howard Rines and Marty Carson for notes and unpublished data related to their *Pc94* introgressions. We thank Roger Caspers for conducting the original mapping population crosses and for general technical assistance along with Jakob Riddle, Krista Ristinen, and Kim-Phuong Nguyen. We thank Aaron Beattie, Melanie Caffe, Nick Metz, Michael McMullen, Sepehr Mohajeri Naraghi, and Kirby Nilsen for sharing pedigree information, germplasm, and unpublished inoculation data.

## Data Availability

The *Avena strigosa* genomes and annotations are available on Grain Genes (https://graingenes.org/GG3/). The GBS and RNAseq data are available in the NCBI SRA database in BioProject PRJNA1381582. All other data supporting the findings of this study are available within the paper, its supplementary information, or on figshare (10.6084/m9.figshare.c.8242072).

