## Supplementary figures 1-7 for "Mapping *Pc94*-Mediated Crown Rust Resistance in Oat Reveals a zfBED NLR and Introgression Variation"

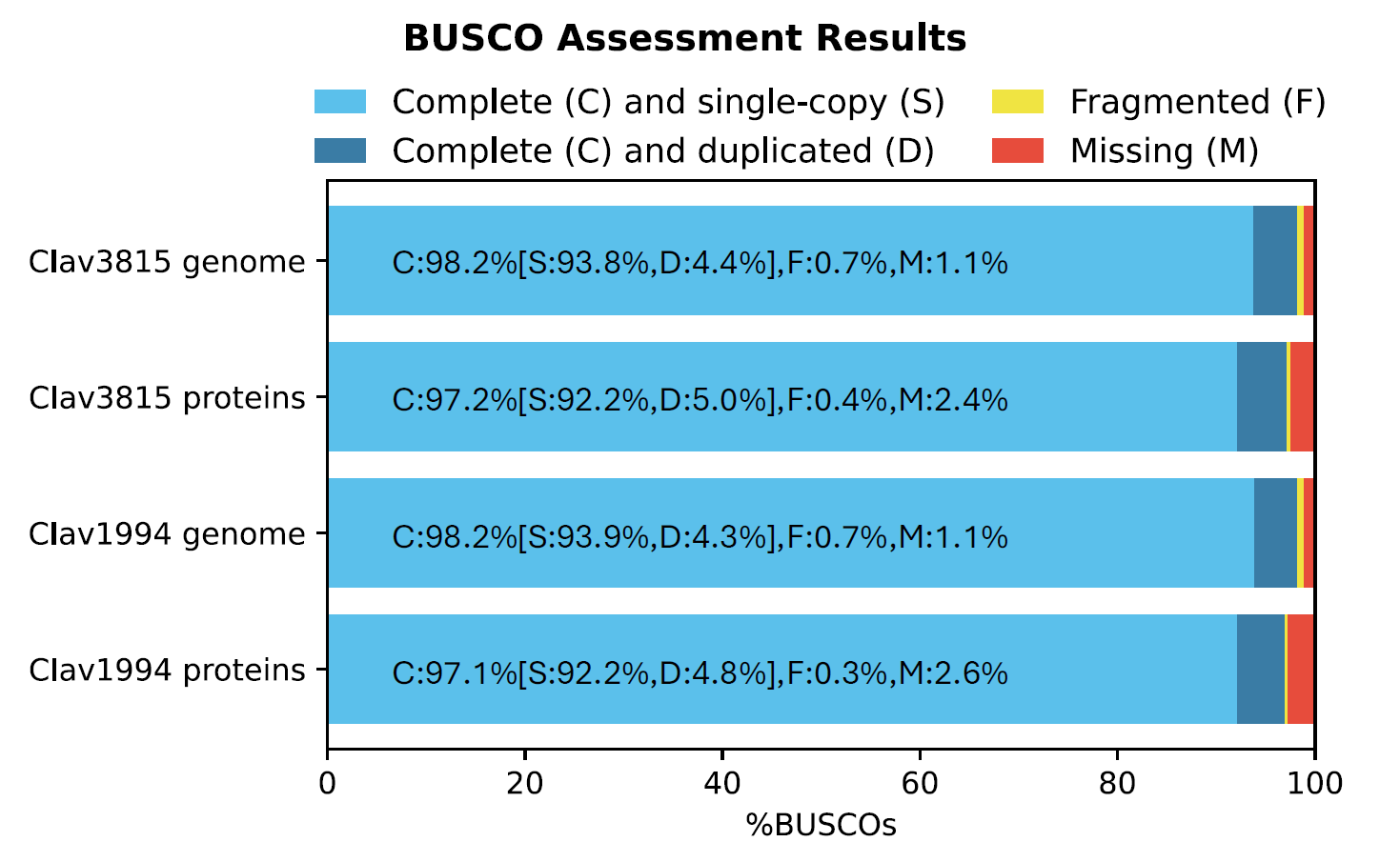

Supplementary figure 1. BUSCO assessment of CIav genome assemblies and proteomes. BUSCO completeness of CIav3815 and CIav1994 genome assemblies and predicted proteomes using the poales_odb10 dataset. Bars represent the percentage of complete single-copy (S), complete duplicated (D), fragmented (F), and missing (M) BUSCOs. Percent values are shown within bars.

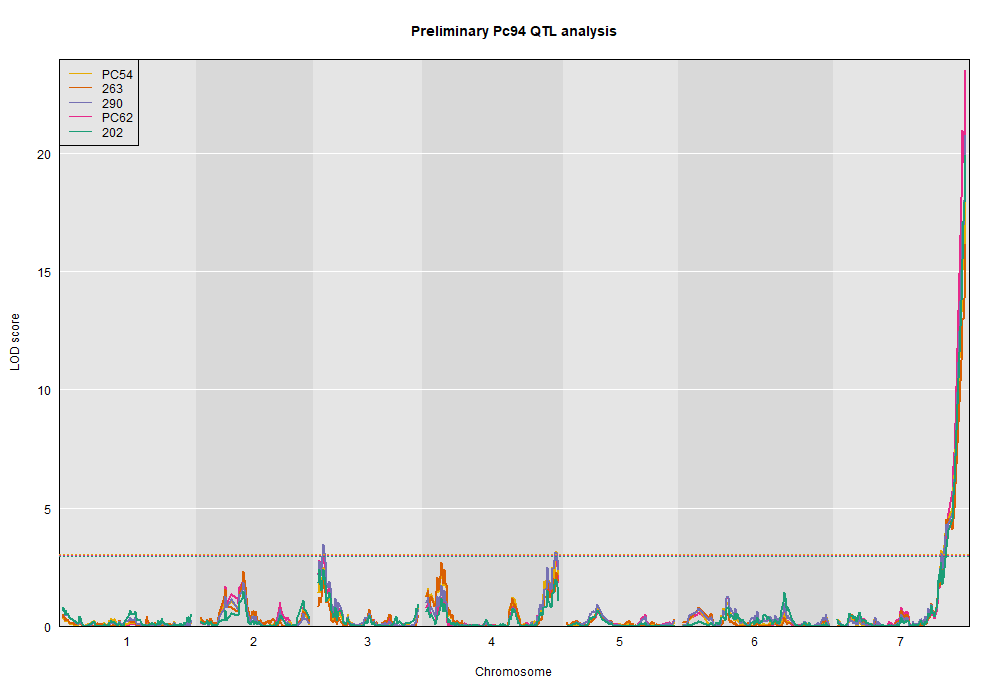

Supplementary figure 2. Preliminary Pc94 QTL analysis plot using previously published linkage maps and phenotypic data for Pca strains PC54, 263, 290, PC62, and 202. The dotted lines are the 0.05 significance thresholds.

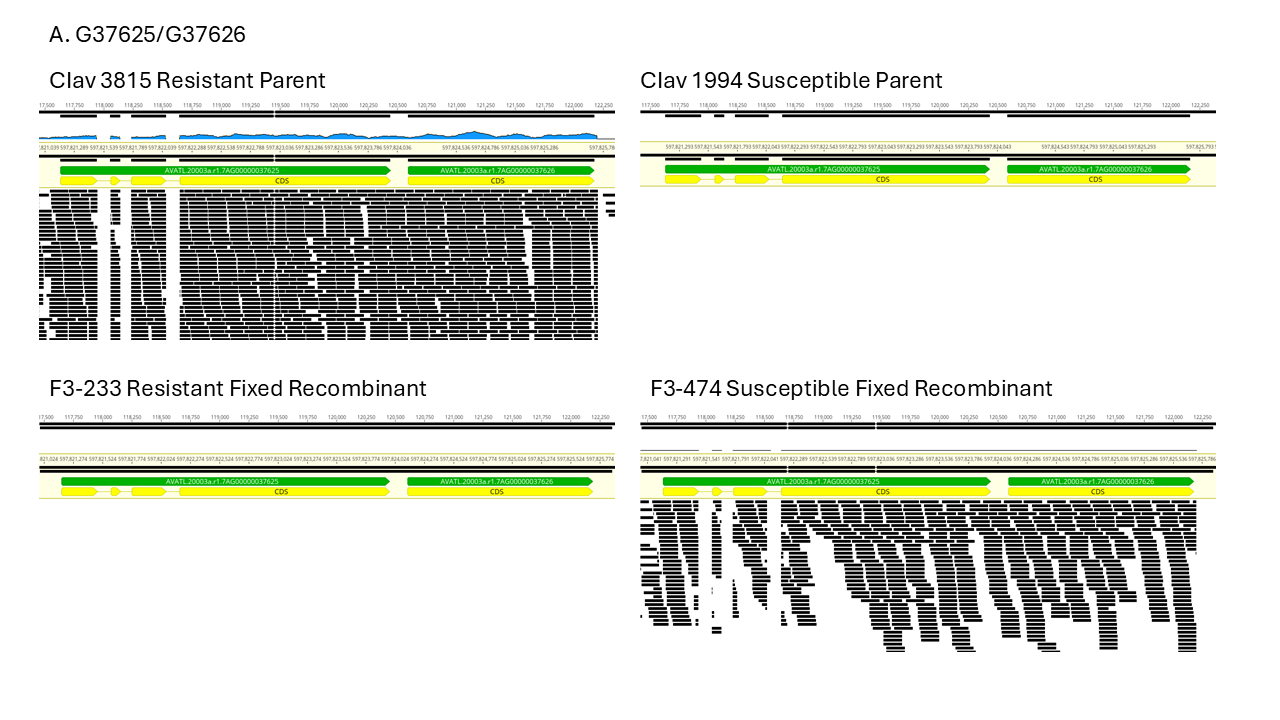

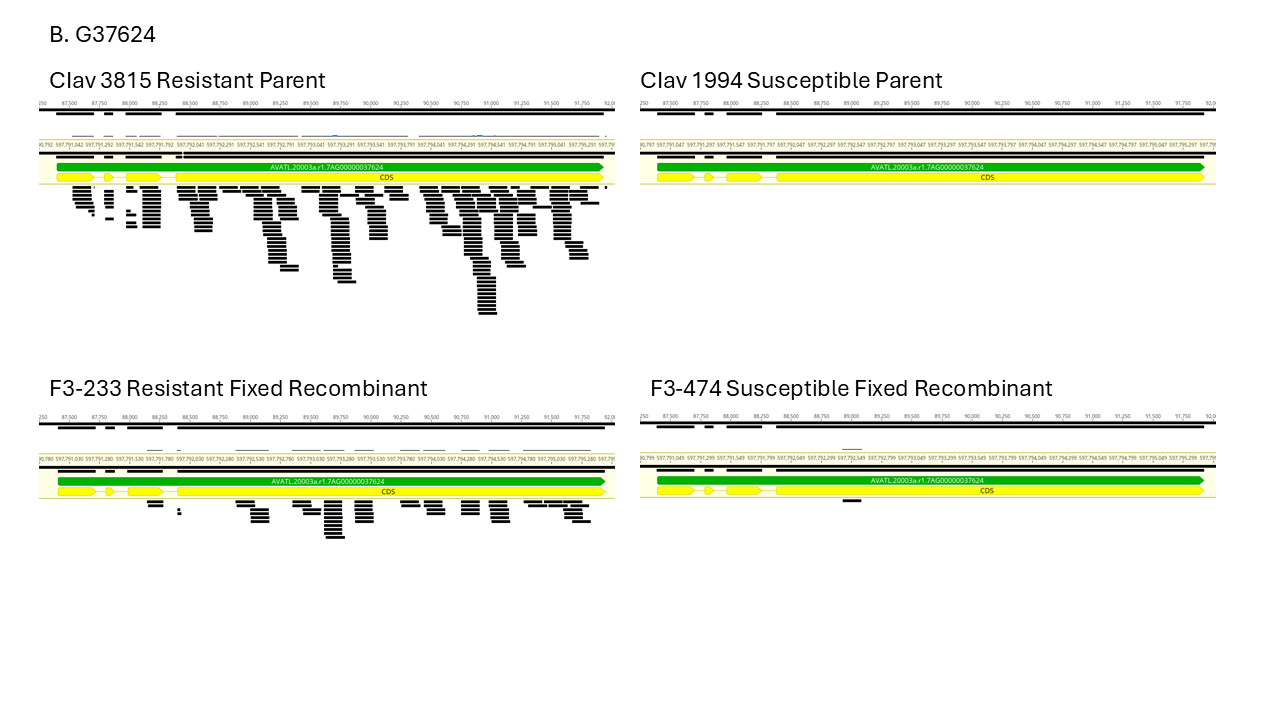

Supplementary figure 3. Leaf tissue RNAseq alignments to NLR genes between KASP markers 7 and 32. A. *G37625/G37626* is not expressed in resistant fixed recombinant F3-233 but expressed in susceptible fixed recombinant F3-474, eliminating it as a candidate gene for *Pc94*. B. *G37624* was much more lightly expressed.

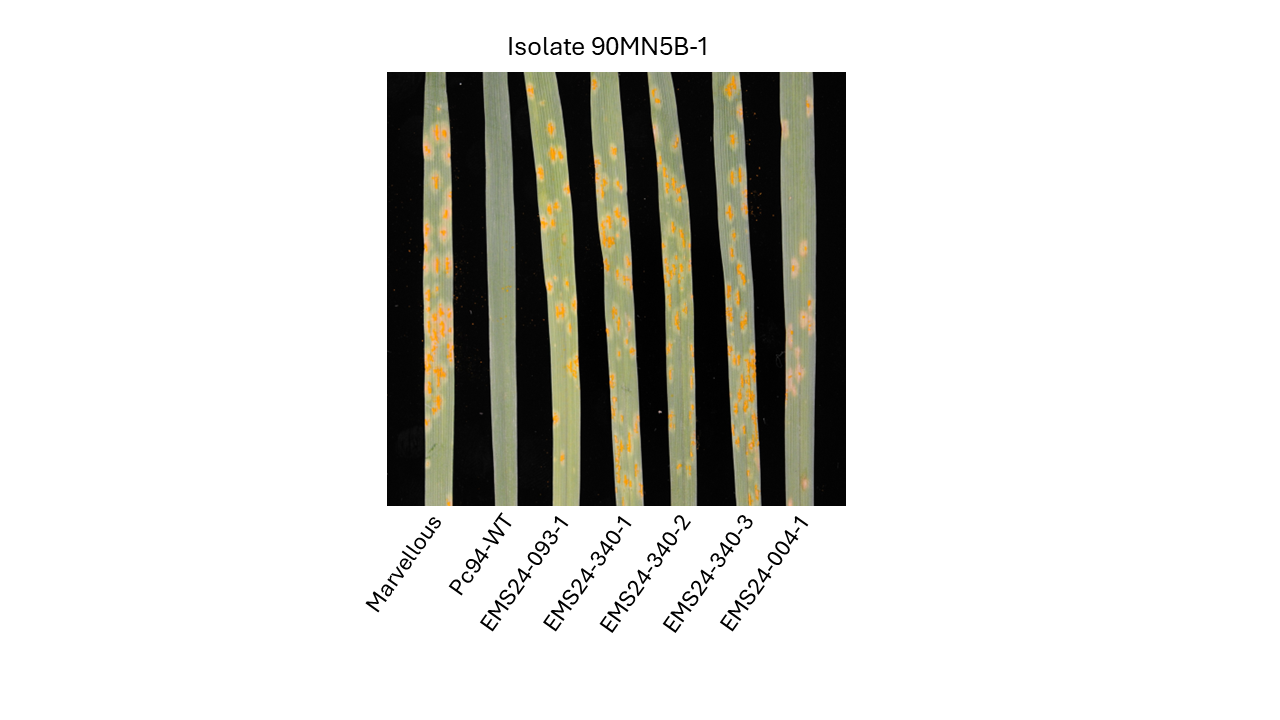

Supplementary figure 4. Phenotypic response of EMS mutants to 90MN5B-1. EMS24-004-1 exhibits a more resistant 2 Infection Type for this isolate compared to the other mutants showing 4 Infection Types.

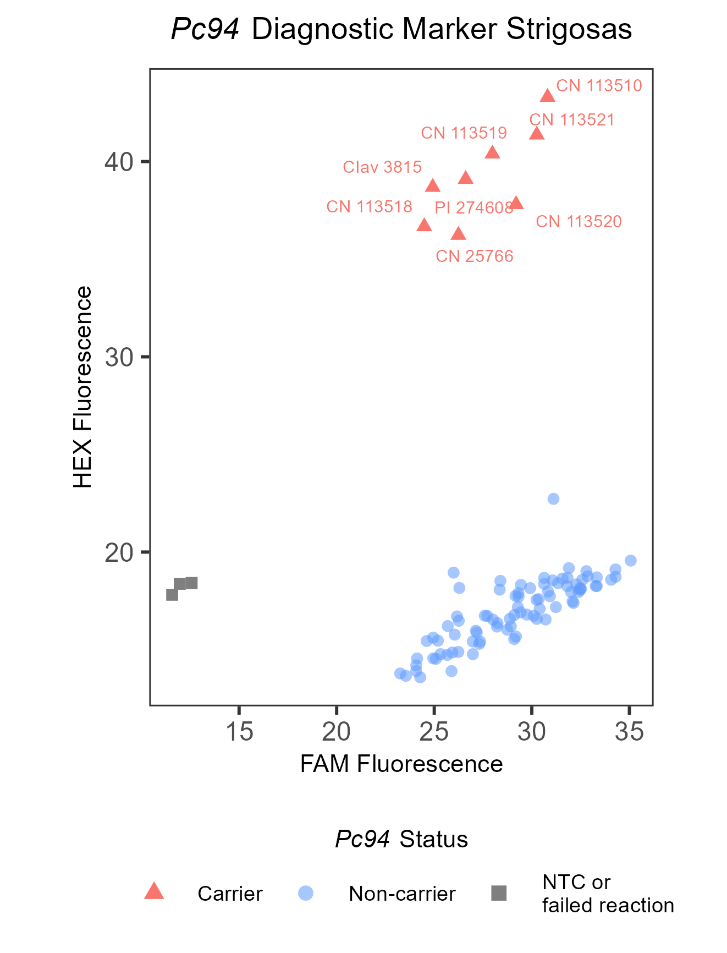

Supplementary figure 5. *Pc94* PACE fluorescence cluster plot. *Avena strigosa* *Pc94* carriers are labeled as red triangles.

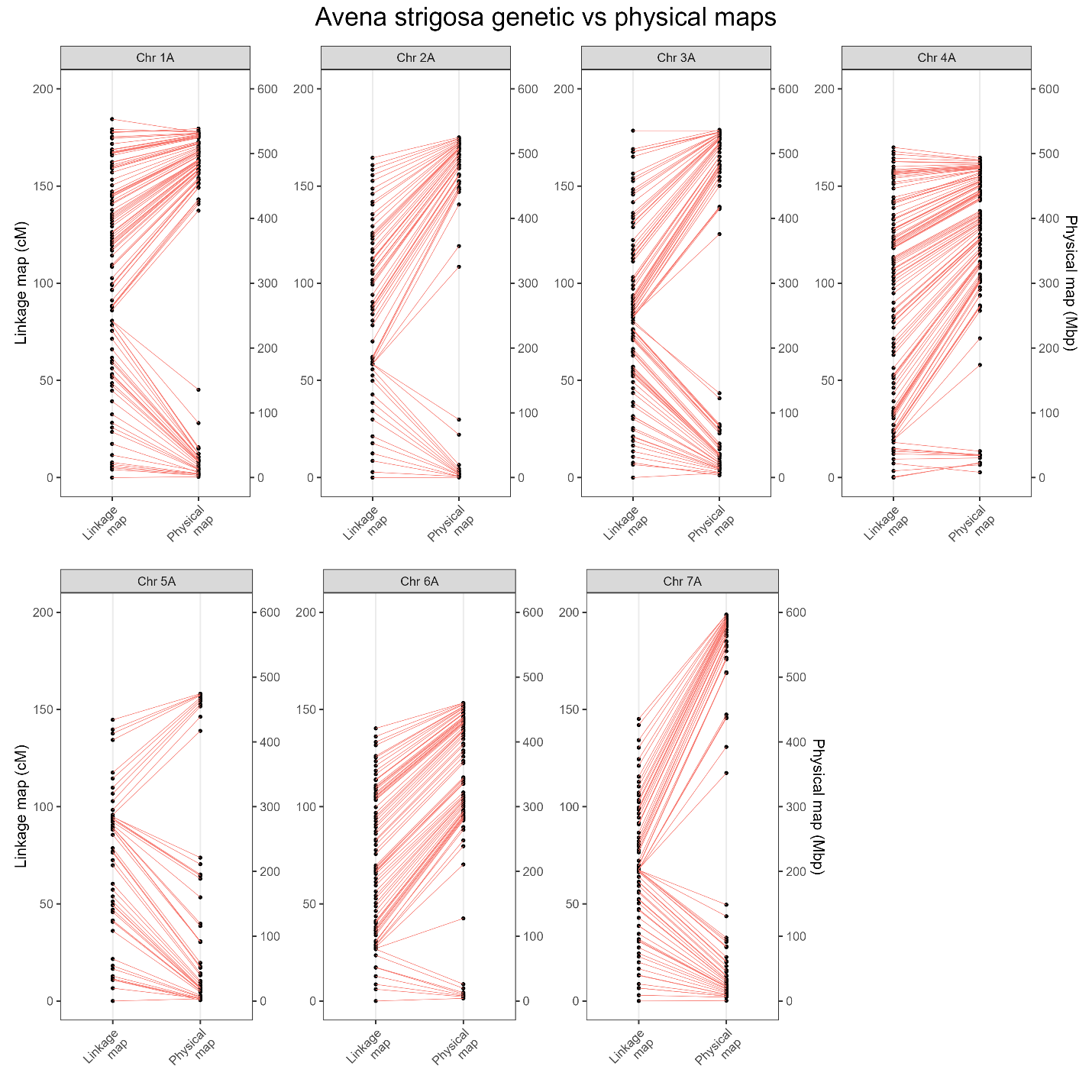

Supplementary figure 6. Comparison of *Avena strigosa* genetic map to the *Avena strigosa* CIav 3815 genome assembly (physical map).

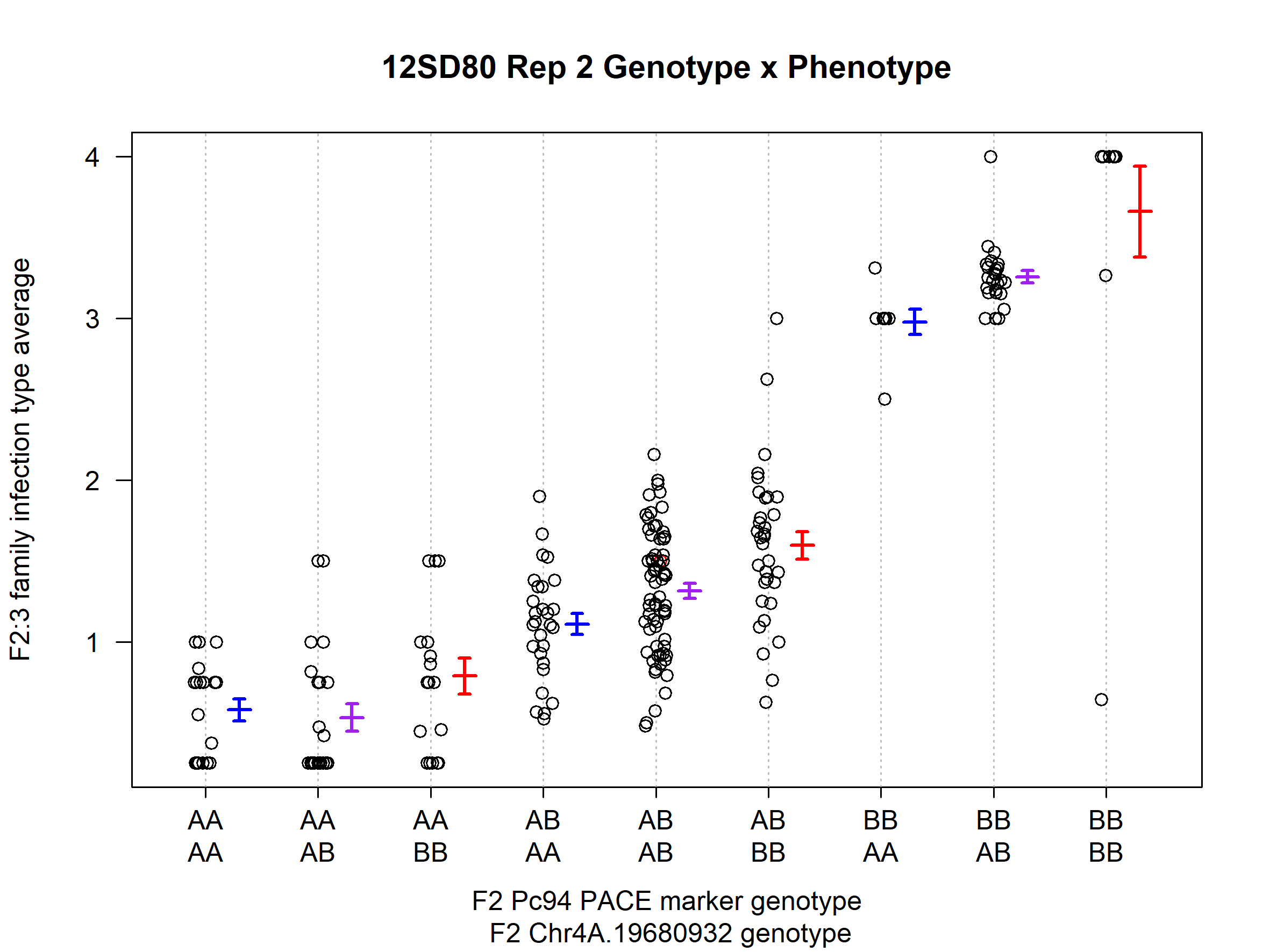

Supplementary figure 7. Genotype x phenotype plot for *Pca* isolate 12SD80 Rep 2. The x-axis denotes the F2 genotypes for the Pc94 PACE marker (first genotype line, QTL on Chr7A) and the Chr4A.19680932 marker (second genotype line, QTL on Chr4A).
